# Multisite Phosphorylation and Binding Alter Conformational Dynamics of the 4E-BP2 Protein

**DOI:** 10.1101/2022.01.14.476386

**Authors:** Spencer Smyth, Zhenfu Zhang, Alaji Bah, Thomas E. Tsangaris, Jennifer Dawson, Julie D. Forman-Kay, Claudiu C. Gradinaru

## Abstract

Intrinsically disordered proteins (IDPs) play critical roles in regulatory protein interactions, but detailed structural/dynamics characterization of their ensembles remain challenging, both in isolation and they form dynamic ‘fuzzy’ complexes. Such is the case for mRNA cap-dependent translation initiation, which is regulated by the interaction of the predominantly folded eukaryotic initiation factor 4E (eIF4E) with the intrinsically disordered eIF4E binding proteins (4E-BPs) in a phosphorylation-dependent manner. Single-molecule Förster resonance energy transfer showed that the conformational changes of 4E-BP2 induced by binding to eIF4E are non-uniform along the sequence; while a central region containing both motifs that bind to eIF4E expands and becomes stiffer, the C-terminal region is less affected. Fluorescence anisotropy decay revealed a nonuniform segmental flexibility around six different labelling sites along the chain. Dynamic quenching of these fluorescent probes by intrinsic aromatic residues measured via fluorescence correlation spectroscopy report on transient intra- and inter-molecular contacts on ns-μs timescales. Upon hyperphosphorylation, which induces folding of ~40 residues in 4E-BP2, the quenching rates decreased at most labelling sites. The chain dynamics around sites in the C-terminal region far away from the two binding motifs significantly increased upon binding to eIF4E, suggesting that this region is also involved in the highly dynamic 4E-BP2:eIF4E complex. Our time-resolved fluorescence data paint a sequence-level rigidity map of three states of 4E-BP2 differing in phosphorylation or binding status and distinguish regions that form contacts with eIF4E. This study adds complementary structural and dynamics information to recent studies of 4E-BP2, and it constitutes an important step towards a mechanistic understanding of this important IDP via integrative modelling.

## INTRODUCTION

Intrinsically disordered proteins (IDPs) are a class of proteins that lack well-defined three-dimensional structures while still carrying out biological functions (1–3). IDPs play a crucial role in mediating interactions with multiple partners and often function as protein interaction hubs (4, 5). IDPs within these protein complexes can undergo disorder-to-order transitions or remain dynamic (6). The lack of stable folded structures observed in IDPs leads to the simplistic assumption that IDPs resemble random coils. In fact, IDPs have transient secondary and tertiary structures and preferential backbone torsion angle propensities due to electrostatic and other interactions based on their amino acid sequence and exhibit a wide range of compactness (6–9).

Cap-dependent initiation of translation is regulated by the interaction of eukaryotic initiation factor 4E (eIF4E) with disordered eIF4E binding proteins (4E-BPs) in a phosphorylation-dependent manner (10–12). The eIF4E protein, which binds the 7-methyl guanosine cap structure of mRNA at the 5’ end, has been shown to be an oncogene and be involved in the induction of cellular transformation (13, 14). The eIF4G, a scaffolding protein, plays a crucial role in docking and assembling several components of the translation initiation machinery at the 5’ cap of mRNA to recruit the ribosome (15). The 4E-BP2 protein is involved in controlling cell growth and proliferation via regulating mRNA translation (16) and in immunity to viral infections (17). Neural 4E-BP2 also functions in regulating synaptic plasticity, playing an essential role in learning and memory, and has been implicated in autism spectrum disorder (18, 19).

The interaction between eIF4E and eIF4G is the fundamental step that initiates the translation process. This interaction involves the canonical binding helix-forming ^54^YXXXXLϕ^60^ (where Y is tyrosine, X is any amino acid, L is Leucine, and ϕ is a hydrophobic amino acid) motif of eIF4G, which is also found in all 4E-BPs, binding to the same convex interface of eIF4E. Thus, the 4E-BPs regulate translation by competing with eIF4G to prevent the assembly of the eIF4F complex and the subsequent mRNA recruitment to the ribosome. Binding of IDPs often leads to ordering, the transient helical structure around the canonical ^54^YXXXXLϕ^60^ motif is stabilized upon eIF4E binding. However, the 4E-BP2:eIF4E complex has been shown by NMR to be highly dynamic with an exchanging bipartite interface (20), in which the secondary binding site ^78^IPGTV^82^ interacts with the lateral surface of eIF4E, as revealed by an X-ray crystal structure capturing a snapshot of the complex (21).

The 4E-BP2 protein is hierarchically phosphorylated. Modification of the first two sites T37 and T46 results in the hypo-phosphorylated state(22); this 2-site phosphorylation (2P) state induces formation of a 4-stranded β-sheet structure from residues 18-62, partially sequestering the canonical binding motif and weakening eIF4E binding but still enabling competition with eIF4G(12). Additional phosphorylation at S65, T70, and S83 leads to a 5-site phosphorylation (5P) state that stabilizes the fold (23), further decreasing the eIF4E affinity and allowing translation initiation to proceed (10). The disordered region C-terminal to the folded domain (C-IDR) remains disordered after phosphorylation and stabilizes the folded domain via long-range interactions (12, 23). However, important details of the structure and dynamics of 4E-BP2 and the eIF4E:4E-BP2 complex remain unknown, which prevent a clear mechanistic picture of the function of 4E-BP2 and its phosphoregulation of translation initiation.

Typically, IDPs have a wide range of interchanging conformations, therefore obtaining dynamic information is critically important for understanding their biological functions. Single-molecule fluorescence (SMF) techniques have been applied previously to measure the conformational heterogeneity, the global dimensions, and the dynamics of IDPs (24, 25). Here we applied a combination of ensemble and single-molecule techniques, i.e., fluorescence anisotropy decay (FAD) and fluorescence correlation spectroscopy (FCS), to characterize global and local peptide chain motions in 4E-BP2 upon multisite phosphorylation and upon binding to eIF4E. FCS and FAD are sensitive to chain motions on the nanosecond-to-microsecond time scale, which is highly relevant for protein folding and for IDP dynamics (26). FAD measurements informed on the local, segmental flexibility of the peptide chain at various sites of the protein, leading to a coarse rigidity map of 4E-BP2 in the non-phosphorylated (NP), 5-site-phosphorylated (5P), and eIF4E-bound states. Using FCS, we resolved up to two different timescales of intra-molecular conformational dynamics in 4E-BP2 under NP, 5P, and denaturing conditions. In addition, we obtained kinetic information (amplitude and lifetime) of key inter-molecular contacts of the dynamic binding interface between NP 4E-BP2 and eIF4E.

Single-molecule Förster resonance energy transfer (smFRET) measurements of two different regions of the protein delineated changes in intramolecular distances and chain rigidity upon multisite phosphorylation, and upon binding to eIF4E. Surprisingly, smFRET showed an increase in the distance between residues 32 and 91 upon phosphorylation, despite folding-induced compaction of residues 18 to 62 (12). While in complex with eIF4E, a region of 4E-BP2 containing both canonical and secondary binding sites expands and stiffens considerably, while the C-terminal region expands only slightly but remains highly flexible. Our multifaceted spectroscopic characterization of the 4E-BP2 conformational dynamics is an important step towards understanding the interplay between folding and release of binding to eIF4E, and it provides valuable information for calculating conformational ensembles of this multistate IDP via integrative modelling.

## RESULTS AND DISCUSSION

### Local chain dynamics measured by FAD

The structural flexibility of IDPs is essential for regulating their interactions with other proteins and their role in signaling processes (3). FAD measures the rotational dynamics of the emission dipole of a fluorophore and is therefore a suitable reporter of the local (segmental) chain flexibility around the labelling site. Inferred parameters from FAD analysis relate to the spatial confinement and the friction experienced by the dye, the movement of a protein segment around the labelling site, and the hydrodynamic radius of the protein segment (27). Probing local conformational dynamics is particularly relevant for IDPs, for which motions of different segments may be uncorrelated and obscured by the averaged global dynamics.

Six single-cysteine 4E-BP2 constructs were used for site-specific fluorescence labelling: C0 (i.e., C0ins/C35S/C73S mutations), C14 (S14C/C35S/C73S), C35 (C73S), C73 (C35S), C91 (C35S/C73S/S91C), and C121 (C35S/C73S/C121ins) (**Fig. S1, Table S1**). While the fastest, sub-ns FAD lifetime (*ρ_dye_*) typically describes the rotation of the dye-linker around the labelling site, in IDPs with transient structure the slowest FAD lifetime describes the rotational diffusion of a protein segment (*ρ_seg_*), where individual segments can rotate independently (28, 29). Alternatively, for folded proteins the slowest FAD lifetime describes the rotational diffusion of the entire protein.(30)

**Fig. 1A** shows a family of FAD curves of NP 4E-BP2 with the fluorophore attached to each of the six different mutated cysteine sites. As dye-protein interactions are expected to be negligible for Atto488 (31) (**Fig. S2**), the variations in the anisotropy decays indicate that the chain flexibility is site/region dependent. FAD fitting parameters using **Eq. 1** (see Methods) are listed in **Table 1**, for each labelling site and each sample condition. **Fig. 1C** shows a comparison between segmental lifetimes (*ρ*_seg_) around each 4E-BP2 labelling site in different phosphorylation states, i.e., NP, 2P, and 5P. These values range from 1.4 ns to 3.4 ns and vary considerably with the labelling site and with the phosphorylation state.

**Figure 1.**
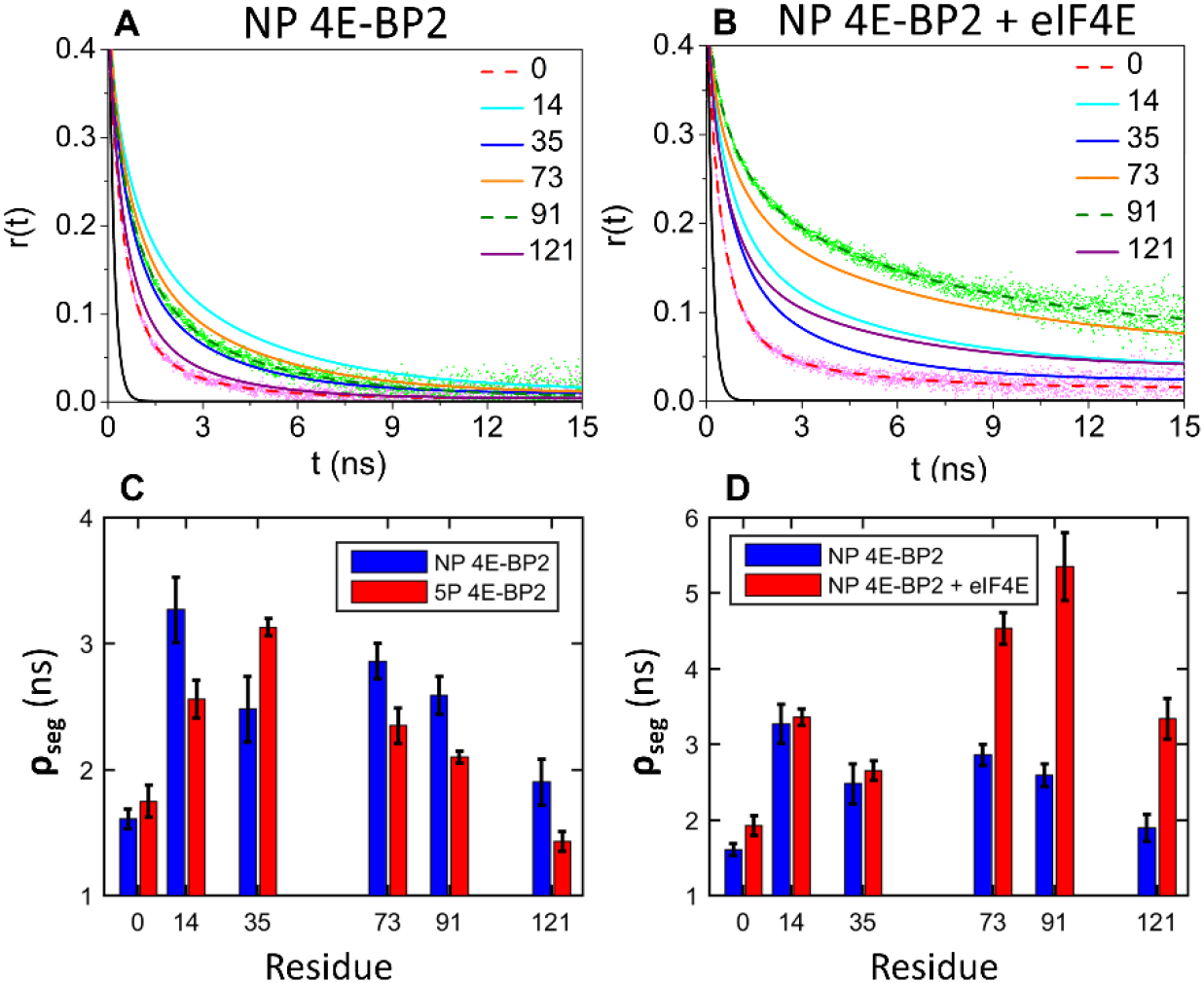
Fluorescence anisotropy decay data for 4E-BP2 fit to **Eq. 1.** The 4E-BP2 protein was labelled with Atto 488 at positions 0, 14, 35, 73, 91 and 121 along the sequence, as described in the text. NP 4E-BP2 fitted curves for all six labelling sites are shown in the absence (**A**) and in the presence of 1 μM of eIF4E (**B**); the FAD of the free dye is shown in black in both panels, exemplar FAD data is shown for positions 0 and 91. (**C**) The slowest rotational lifetime (*ρ_seg_*) obtained by fitting FAD curves at each labelling site for NP (**blue**) and 5P (**red**) 4E-BP2. (**D**) The slowest rotational lifetime (*ρ_seg_*) obtained by fitting FAD curves at each labelling site for NP 4E-BP2 in the absence (**blue**) and in the presence of (**red**) of eIF4E. The concentration of 4E-BP2 used in these experiments was ~50 nM.

**Table 1.** Anisotropy decay parameters for phosphorylated/bound 4E-BP2 states labelled at different sites ^a^.

|  | **5P 4E-BP2** |  | **NP 4E-BP2** |  | **NP 4E-BP2 + elF4E** |  |
| --- | --- | --- | --- | --- | --- | --- |
| | $\rho_{\text{dye}}$ (ns) | $\rho_{\text{seg}}$ (ns)<br>(a) | $\rho_{\text{dye}}$ (ns) | $\rho_{\text{seg}}$ (ns)<br>(a) | $\rho_{\text{dye}}$ (ns) | $\rho_{\text{seg}}$ (ns)<br>(a) |
| **C0** | 0.53 | 1.75<br>(0.72) | 0.51 | 1.61<br>(0.66) | 0.52 | 1.93<br>(0.70) |
| **S14C** | 0.56 | 2.56<br>(0.51) | 0.64 | 3.27<br>(0.42) | 0.62 | 3.36<br>(0.49) |
| **C35** | 0.55 | 3.13<br>(0.41) | 0.64 | 2.48<br>(0.52) | 0.62 | 2.65<br>(0.51) |
| **C73** | 0.55 | 2.35<br>(0.54) | 0.62 | 2.86<br>(0.47) | 0.61 | 4.54<br>(0.46) |
| **S91C** | 0.57 | 2.01<br>(0.51) | 0.61 | 2.59<br>(0.50) | 0.65 | 5.35<br>(0.43) |
| **C121** | 0.52 | 1.43<br>(0.61) | 0.53 | 1.90<br>(0.60) | 0.59 | 3.34<br>(0.59) |

<sup>a</sup> All data were fit to **Eq. 1** to estimate two rotational correlation lifetimes (dye & segment) and their fractions (a is the fraction of the slowest component). Fitting error margins are on the order of $\pm 0.1$ ns for the lifetimes and $\pm 0.05$ for amplitudes (**Table S2**).

Many IDPs are more compact than an ideal statistical coil of the same length due to transient intra-molecular contacts (7, 8, 32). Slower segmental dynamics in PBS buffer (pH 7.4, 140 mM NaCl) were observed than in chemical denaturant (6M GdmCl) at all labelling sites (**Table S2**). The chemically denatured NP and 5P states of 4E-BP2 have similar segmental flexibility signatures, with *ρ_seg_* = 0.8 – 1.0 ns at the ends and *ρ_seg_* = 1.3 – 1.6 ns at internal sites. The values and the trend here match previous measurements for denatured proteins and the expectations for a random coil state(27, 29). Additionally, the pattern of FAD lifetimes is consistent overall with that of ^15^N relaxation rates(20) and of ^1^H-^15^N nuclear Overhauser effect (NOE) values (12). This suggests that FAD-measured segmental dynamics can probe transient secondary structures and non-random chain contacts, and reports on the local degree of disorder in IDPs.

Another contribution to the heterogeneous segmental flexibility is the amino acid composition around each labelling site, with glycine and serine being the most flexible, and proline, isoleucine, and valine the most rigid (33). With 4 serine and 2 glycine among the first 10 residues, the N-terminus is the most flexible region of NP 4E-BP2. Considering a 10-residue window, the positions 14, 73 and 91 are flanked by several rigid residues (proline, isoleucine, valine) and the protein appears to be much less flexible in these regions. The slowest rotational lifetime was observed at position 14 (*ρ_seg_* = 3.27 ns), which, in addition to two proline residues, has two positively charged arginine residues in its proximity, which may rigidify the segment further via electrostatic interactions.

In previous NMR studies, we have shown that a region of 4E-BP2 (residues 18 to 62) folds upon phosphorylation, while the rest of the chain remains disordered (12). We used FAD to probe the changes in segmental flexibility of 4E-BP2 that are induced by complete (5P) phosphorylation (**Fig. 1C, Table 1**). The rotational lifetime (*ρ_seg_*) increases at position 35 while it decreases at all the other dye positions, which are outside the folded domain, indicating that the chain becomes more flexible. This is consistent with the formation of the four-stranded beta-sheet fold between residues 18-62 and with the C-terminal region remaining disordered after phosphorylation. From an entropic perspective, increased conformational flexibility near the secondary binding site (residues 78 to 82) may also contribute to decreasing the affinity for eIF4E.

FAD has been used previously to quantify and differentiate local binding constants of IDPs in the context of multisite interactions (28). **Fig. 1B** shows anisotropy decay data and fitted curves for the NP 4E-BP2:eIF4E complex at each of the six labeling sites on 4E-BP2. In contrast with the apo sample, these curves decay to significantly higher asymptotic values (*r_inf_*, **Table S2**), indicating that the local motions around each labeling site in 4E-BP2 are hindered after binding to eIF4E. The largest changes in chain flexibility occur at three C-terminal sites while the changes observed in the N-terminal sites are minor (**Fig. 1D**). At positions 73, 91 and 121, the segmental rotational lifetime *ρ_seg_* nearly doubles, from 2-3 ns in the apo state to 4-5 ns in the bound state. Similarly, *r_inf_* values significantly increase, with the largest changes at the C-terminal labelling sites (**Fig. S3**).

Positions 73 and 91 are located near the secondary binding site while position 121 is far from either binding site but the changes in lifetime are consistent with binding-induced changes to NMR intensity ratios(20), which demonstrate complete broadening for residues 45-88, significant broadening for residues 34-90, and broadening at residue 120. The data for positions 73 and 91 likely reflect favorable interactions at the secondary binding site, as well as potentially the competitive interaction of the secondary binding region with the disordered N-terminus of eIF4E that has been suggested to act as a negative regulator (34). The NMR broadening results for position 120 together with these FAD data on position 121 support a picture of the 4E-BP2:eIF4E dynamic complex involving a more extensive part of the C-terminus.

The results demonstrate that segmental motion parameters measured by FAD can be used to disentangle the binding contributions of different regions in an intrinsically disordered protein as it interacts with its binding partners. More specifically, the changes in *r_inf_* are correlated with the changes in *ρ_seg_*, indicating that the rotational freedom of the segment and the segmental dynamics of 4E-BP2 are hindered in the bound state.

### Non-local chain dynamics in 4E-BP2 measured by PET-FCS

The same Atto488-labelled 4E-BP2 constructs used for FAD (**Fig. S1, Table S1**) were also used for FCS experiments. FCS is sensitive to intensity fluctuations caused by the diffusion of the labelled protein and by the photophysics of the fluorophore (**Eq. 2**). Notably, the fluorophore can be dynamically quenched by aromatic amino acids via photoinduced electron transfer (PET)(35, 36). Tryptophan and tyrosine are the strongest PET quenchers, with a quenching range of 5-10 Å for typical fluorophores (37, 38). The two tyrosines in the 4E-BP2 sequence (Y34 and Y54) are most likely acting as quenchers in our FCS measurements. Given that the PET quenching effect in our measurements is not produced by a single residue, we do not employ a specific kinetic model and instead limit our interpretation to the PET induced exponential lifetimes and amplitudes accounting for in our fitting model (**Eq. 2**).

**Fig. 2A** shows experimental FCS data, fitting curves, and residuals for a representative sample, 4E-BP2 labelled with Atto488 at position 73, in the NP 4E-BP2:eIF4E state. Models with one diffusion component and two or three (faster) kinetic components (**Eq. 2**) satisfactorily fit the experimental autocorrelation decays measured for all 4E-BP2 samples. The (sub-diffusion) kinetic components in FCS data are attributed to intrinsic triplet-state kinetics of the probe and to dynamic PET quenching induced by the protein environment.

**Figure 2.**
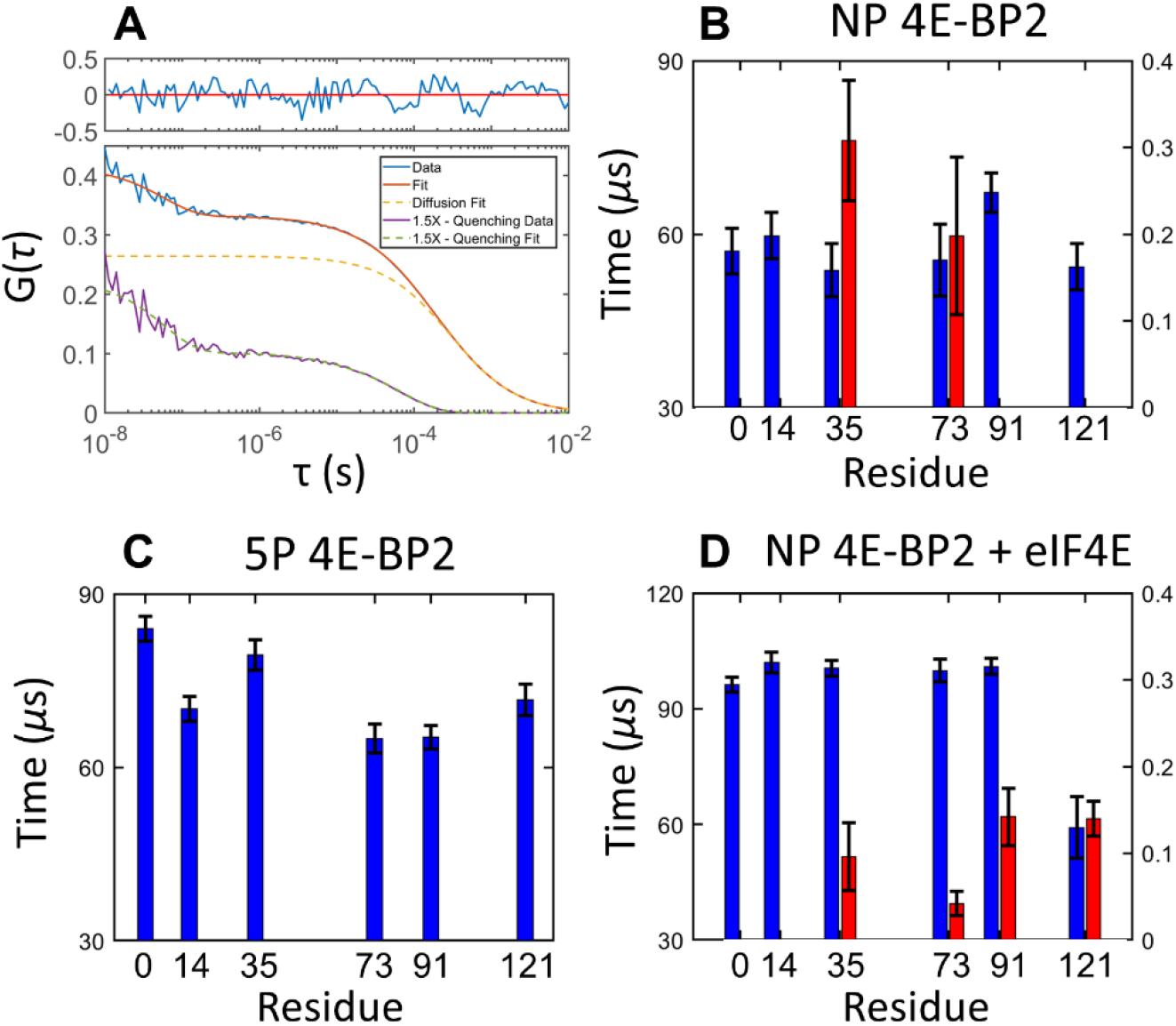
FCS lifetimes of 4E-BP2 labelled with Atto488 at residue 0, 14, 35, 73, 91 or 121. Experimental curves were fit to **Eq. 2**. Fitting of NP 4E-BP2 labeled at residue 73 in the presence of 0.5 μM eIF4E, the kinetic decays and diffusion for the best fit plotted separately (**A**). FCS experiments were performed on 4E-BP2 at a concentration of ~5 nM in different states: non-phosphorylated (**B**), hyper-phosphorylated (**C**), in the presence of 0.5 μM of elF4E (**D**). Different decay timescales are shown in different colors: τ_1_≈100 μs (**blue**) and τ_2_=0.05-0.3 μs (**red**). The fitting error bars for each lifetime are shown in the figure. The full list of fitting parameters is given in **Table S3**.

The FCS lifetimes obtained by fitting the data to **Eq. 2** can be grouped into three timescales: **τ_1_** ≈ 50-100 μs, **τ_t_** ≈ 7-8 μs, and **τ_2_** ≈ 50-300 ns, respectively (**Fig. 2, Table 2, Fig. S4**). Conversely, the free dye exhibits a single kinetic component with a lifetime of ~3-10 μs (**Fig. S5**), which is similar to a previously reported triplet lifetime of Atto488.(39) Thus, for 4E-BP2, the **τ_1_** and **τ_2_** lifetimes are attributed to PET quenching of the dye due to conformational dynamics of the protein, while the **τ_t_** lifetime was assigned to the intrinsic triplet lifetime of Atto488. Therefore the (**τ_t_**) and (**a_t_**) component were fit as shared parameters in each measurement condition, the diffusion coefficient is also a shared (see methods). Chain motions on the sub-μs (**τ_2_**) timescale are typically associated with interconversion of states within the disordered conformational ensemble and with proximal loop formation.(40, 41). The slowest (**τ_1_**) kinetics are on the same timescale as concerted motions of the protein chain, such as domain movements or transient tertiary structural contacts (43, 44).

**Table 2.**
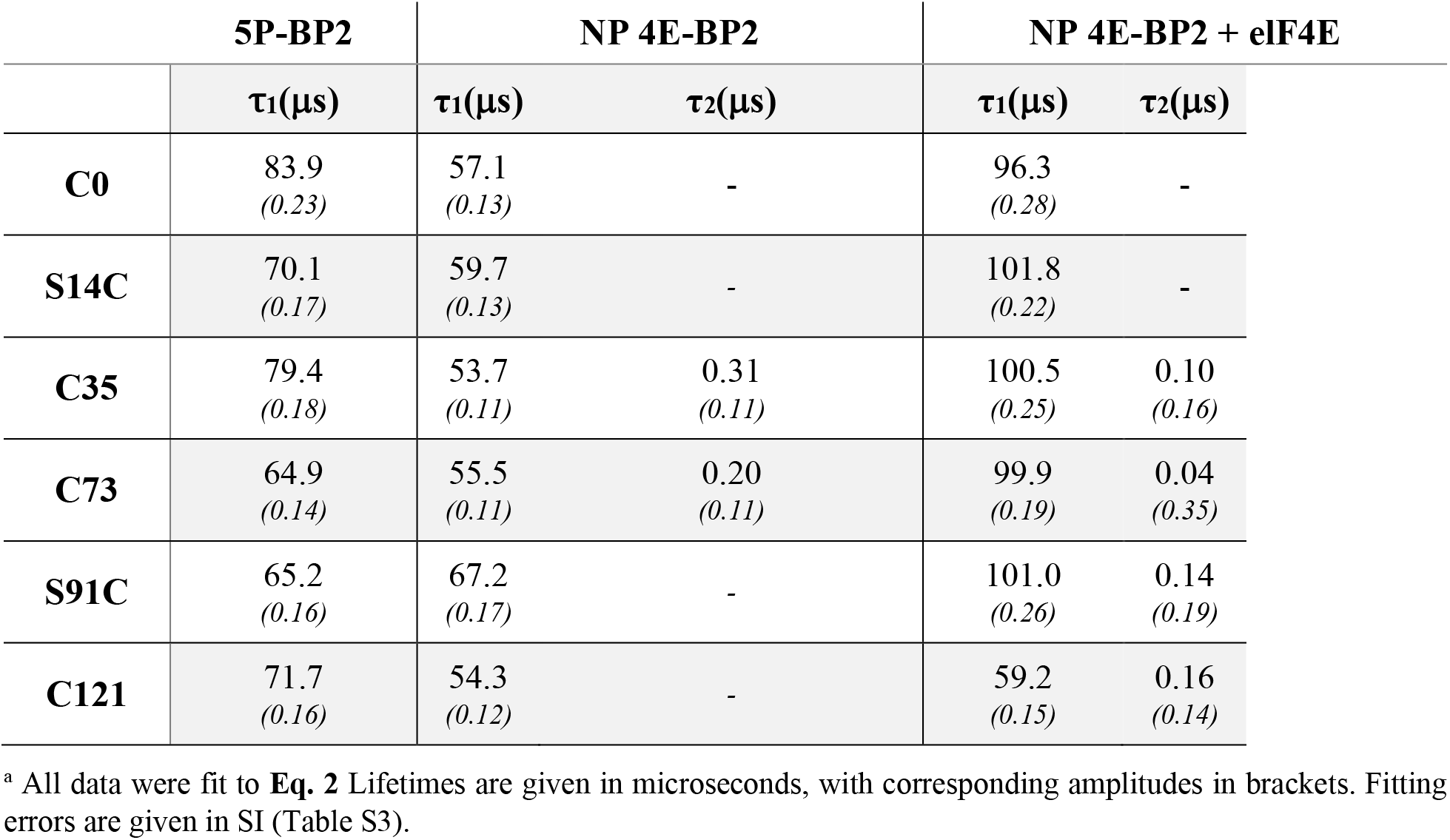
FCS decay parameters for five-phospho, non-phospho and elF4E-bound 4E-BP2 ^a^.

|  | 5P-BP2 | NP 4E-BP2 |  | NP 4E-BP2 + eIF4E |  |
| --- | --- | --- | --- | --- | --- |
| | $\tau_1(\mu\text{s})$ | $\tau_1(\mu\text{s})$ | $\tau_2(\mu\text{s})$ | $\tau_1(\mu\text{s})$ | $\tau_2(\mu\text{s})$ |
| **C0** | 83.9<br>(0.23) | 57.1<br>(0.13) | - | 96.3<br>(0.28) | - |
| **S14C** | 70.1<br>(0.17) | 59.7<br>(0.13) | - | 101.8<br>(0.22) | - |
| **C35** | 79.4<br>(0.18) | 53.7<br>(0.11) | 0.31<br>(0.11) | 100.5<br>(0.25) | 0.10<br>(0.16) |
| **C73** | 64.9<br>(0.14) | 55.5<br>(0.11) | 0.20<br>(0.11) | 99.9<br>(0.19) | 0.04<br>(0.35) |
| **S91C** | 65.2<br>(0.16) | 67.2<br>(0.17) | - | 101.0<br>(0.26) | 0.14<br>(0.19) |
| **C121** | 71.7<br>(0.16) | 54.3<br>(0.12) | - | 59.2<br>(0.15) | 0.16<br>(0.14) |

<sup>a</sup> All data were fit to **Eq. 2** Lifetimes are given in microseconds, with corresponding amplitudes in brackets. Fitting errors are given in SI (Table S3).

For NP 4E-BP2, one or two kinetic components were resolved in addition to the triplet component at each labelling position, which follow the **τ_1_** and **τ_2_** timescales. The lifetimes for the **τ_1_** components are site-dependent (**Fig. 2B**). The fastest component **τ_2_** was only resolved at positions C73 and S91C, this component was not resolved for the other positions within experimental error limits. Similar PET-FCS analysis, with multiple lifetimes of intrachain dynamics between ~100 ns and ~100 μs, was previously reported for other IDPs, e.g., the N-terminal domain of p53-TAD (42) and the mouse prion protein moPrP (43).

In contrast to the NP state, the 5P state exhibits only one FCS kinetic component (**τ_1_**) in addition to the triplet component at each labelling position (**Fig. 2C**). In the 5P state, the addition of phosphate groups and the induced folding in the 18-62 region probably slows down the protein reconfiguration time sufficiently to be “absorbed” into the low μs triplet dynamics, as previously reported for p53-TAD (42). Upon phosphorylation, the **τ_1_** component slows down at all positions except 73 and 91 where no change is observed within the experiment error limits; the associated kinetic amplitude ***a*_1_** decreases at all sites except for position 91 (**Table 2, Fig. S6**), from which it can be inferred that the dominant effect of phosphorylation is to slow down intrachain contacts causing PET quenching. Similarly, slower conformational dynamics upon multisite phosphorylation has been observed for the disordered p53-TAD protein, even though phosphorylation does not lead to folding in this system (42). In 4E-BP2, phosphorylation stabilizes the β-strand structure between residues 18 and 62, which is expected to limit the access of the fluorophore to tyrosine quenchers (12).

Given the tight binding affinity of 4E-BP2 for eIF4E (*k_d_* = 3.2 ± 0.6 nM)(12), saturating amounts of eIF4E (0.5 μM) were used to ensure that nearly all (>99%) 4E-BP2 molecules were in the bound state. FCS measurements on Atto488-labelled 4E-BP2 with or without unlabeled eIF4E (**Fig. S2**) showed a shift to longer diffusion times (larger *R_H_*) in the presence of eIF4E, which is consistent with the formation of the 4E-BP2:eIF4E complex. The *R_H_* values estimated by fitting the data to **Eq. 2** are 29.0 ± 0.5 Å for the NP 4E-BP2 (independent of labelling site) and 36.6 ± 0.8 Å for the 4E-BP2:eIF4E complex. As reference, FCS measurements of Atto488 (5 nM) in the presence of excess eIF4E (1 μM) showed that the dye does not interact/bind with eIF4E (**Fig S2**).

When bound to elF4E, NP 4E-BP2 exhibits one to two FCS kinetic components in addition to the triplet component which fall into the same τ_1_ and τ_2_ range as in the free (Apo) state, (**Fig. 2D**). eIF4E (PDB: 3AM7) contains 8 tryptophan and 6 tyrosine residues, of which four tryptophan (W46, W56, W73 and W102) and three tyrosine (Y34, Y76, and Y 145) residues are largely surface exposed (**Table S4**). Therefore, the observed changes in PET quenching dynamics are the combined result of the new inter-molecular interactions between 4E-BP2 and the surface of eIF4E and the changes in intra-molecular contacts within 4E-BP2.

4E-BP2 interacts with eIF4E through dynamic interactions involving at least an α-helical structure at the canonical ^54^YXXXXLϕ^60^ motif and the secondary ^78^IPGVT^82^ site (20, 21). Positions 35 and 73 show a decrease in lifetime accompanied by an increase in amplitude of the fast component, positions 91 and 121 gain this component upon binding to eIF4E (**Fig. 3 C-F**). As such, τ_2_ decreases from ~200-300 ns to ~50-100 ns upon binding at positions 35 and 73, while τ_2_ at positions 91 and 121 is ~150 ns when bound (**Table 2**). The appearance of the fast τ_2_ component at positions 91 and 121, as well as the decrease in τ_2_ lifetime and increase in *a*_2_ amplitude at position 73 are likely caused by dynamic exchange of the secondary binding site leading to quenching by aromatic residues on the surface of eIF4E. The involvement of residues 73, 91 and 121 in eIF4E binding is consistent with binding-induced NMR intensity changes reported previously(20). The changes at residue 121 upon binding mirror those observed by FAD (**Fig. 1 D**), providing further support for a dynamic complex involving more of the C-terminal portion of 4E-BP2. Taken together, the results highlight the dynamic nature and range of timescales present in the bound state and ability of PET-FCS to probe intermolecular dynamics within the bound state.

**Figure 3.**
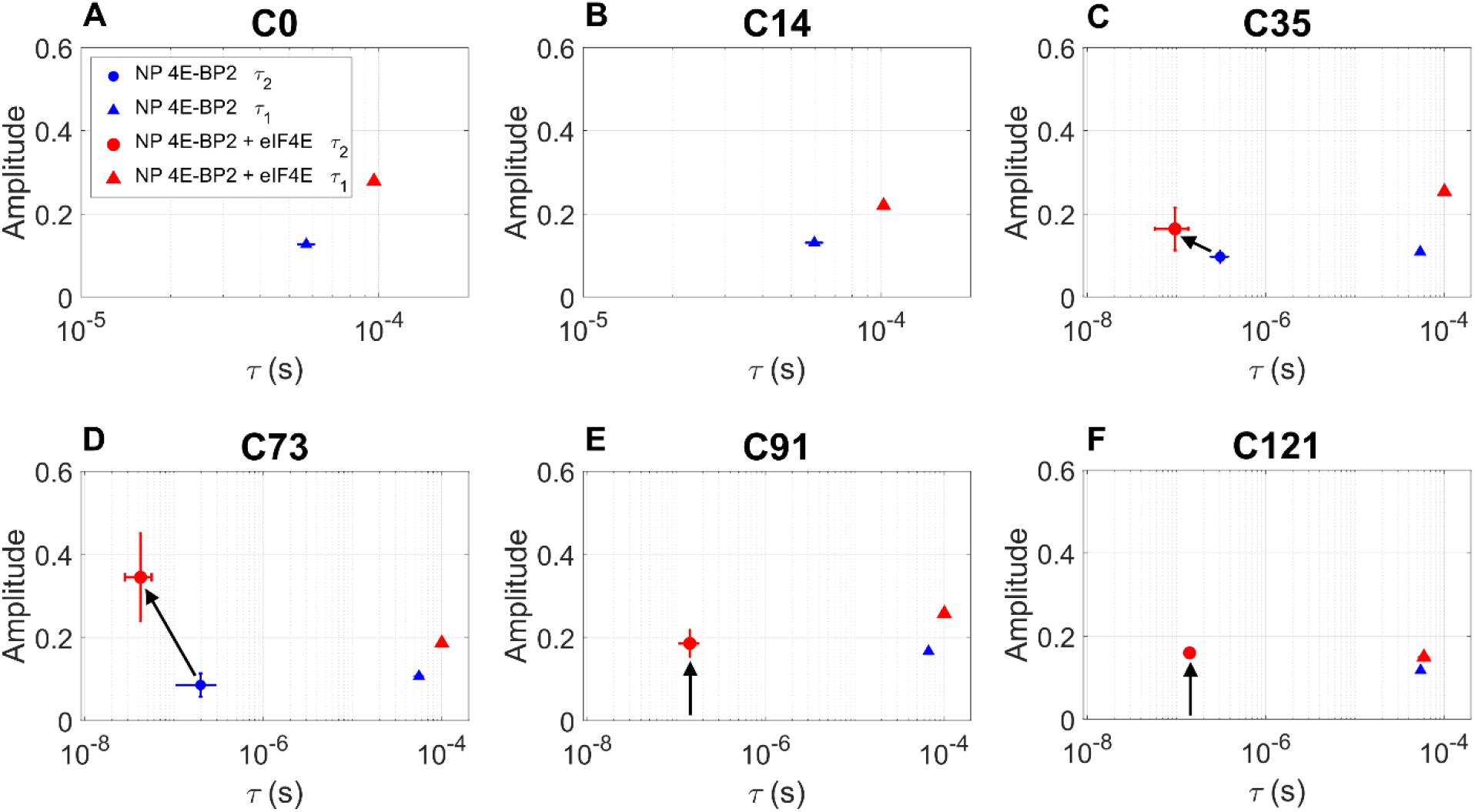
The changes in lifetime and amplitude of the fastest and slowest FCS kinetic components (**τ_2_ – circle**, and **τ_1_ – triangle**) induced by binding of NP 4E-BP2 to elF4E. FCS measurements on NP 4E-BP2 labelled with Atto488 at six different sites were performed in the Apo state (**blue**) and in the presence of 0.5 μM eIF4E (**red**), and the data were fit to **Eq. 2.** The fitting error bars for each parameter are indicated in the figure; some error bars are too small to be seen, all fitting parameters are given in **Table S3**.

### Chain dimensions and stiffness of 4E-BP2 assessed by smFRET

smFRET is exquisitely suited to delineate heterogeneous and dynamic states that are inherent to IDPs. smFRET can resolve heterogeneous population distributions and kinetics, is compatible with a wide range of solution conditions, and overcomes the ensemble averaging of established structural techniques such as NMR and small-angle X-ray scattering (SAXS)(25). By recording the arrival time, the color and the polarization of each detected photon, multiparameter fluorescence (MPF) detection permits access to additional intrinsic properties of fluorescence that can be related to properties of the conjugated molecule (46). The chain dimensions and stiffness of 4E-BP2 were assessed using smFRET with MPF on two double-cysteine FRET constructs, which were labelled stochastically with Alexa Fluor 488 (donor) and Alexa Fluor 647 (acceptor).

smFRET histograms for H32C/S91C in the NP, 5P, and NP+eIF4E states are shown in **Fig. 4**. Although each histogram was satisfactorily fit to a single Gaussian, a single FRET peak does not necessarily reflect a homogenous distribution of states. Indeed, the underlying population is likely in fast exchange compared to the burst duration (~1 ms), given the typical chain reconfiguration times of IDPs (~100 ns) (51). The data suggests that the NP state is more compact than a statistical coil ensemble, i.e., 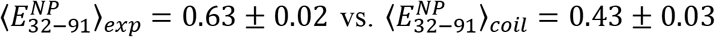 (see SI section 2.3). This is consistent with the presence of transient secondary structure observed by NMR(20), and with the multiple transient intrachain contacts observed by PET-FCS.

**Figure 4.**
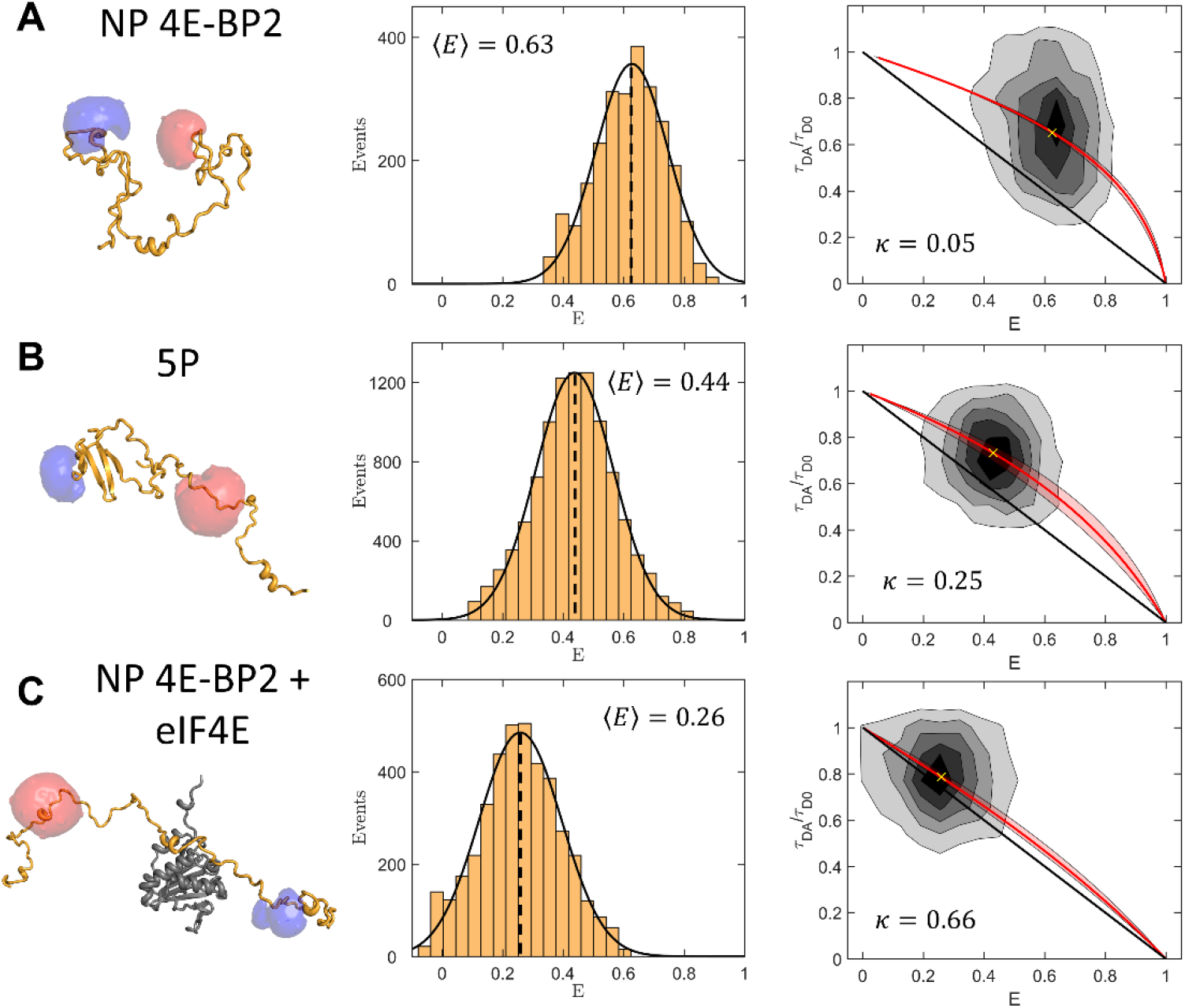
smFRET MPF results for the H32C/S91C 4E-BP2 construct labelled with Alexa Fluor 488 and Alexa Fluor 647 in the (**A**) non-phospho (NP), (**B**) five-phospho and (5P) (**C**) non-phospho eIF4E-bound (NP 4E-BP2 + eIF4E) states. (1^st^ column) Cartoon depictions of conformations of 4E-BP2 in different states with representations of fluorophore accessible volume simulations (47). (2^nd^ column) FRET efficiency histograms fit to a Gaussian function, with the dashed lines indicating the mean values. (3^rd^ column) 2D histogram plots of the FRET efficiency (*E*) and the donor-only normalized fluorescence lifetime; τ_DA_ and τ_D0_ are donor lifetimes in the presence and absence of acceptor, respectively. The black line shows the relation expected for a static rigid molecule. The red line is a relation for a worm-like chain with a stiffness parameter (κ) that passes through the centroid of a 2D gaussian fit indicated by a yellow cross. The red shaded region shows the uncertainty range of κ.

Upon multi-site phosphorylation 4E-BP2 undergoes folding to a beta-sheet domain between residues 18 and 62 (12). Typically, when proteins fold, FRET efficiency increases, following the overall compaction of the structure and contraction of most (but not all!) inter-residue distances(48, 49). In this case however, the FRET efficiency of the 5P state is lower for the dye pairs at residues 32 and 91, 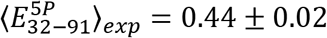, not higher than the NP state, 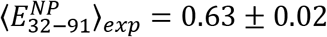 (**Table 3**). The dye at C32 is positioned in the long loop between strands β1 and β2 which is proximal to the C-terminal end of the domain (PBD ID 2MX4), from where the C-IDR, containing the dye at C91, extends. An alternate construct designed to specifically probe the folded region (with both dyes within the folded domain boundaries from 18-62) would likely lead to an increase in FRET efficiency upon phosphorylation. The H32C/S91C construct, however, also partially probes the C-IDR, which expands when 4E-BP2 is phosphorylated (**Fig. 5B**).

**Table 3.** Mean smFRET efficiency and WLC stiffness values of the non-phospho (NP), five-phospho (5P) and elF4E-bound 4E-BP2 (NP 4E-BP2 + eIF4E).

|  | NP 4E-BP2 |  | 5P-BP2 |  | NP 4E-BP2 + eIF4E |  |
| --- | --- | --- | --- | --- | --- | --- |
| | $\langle E \rangle$ | $\kappa$ | $\langle E \rangle$ | $\kappa$ | $\langle E \rangle$ | $\kappa$ |
| **H32C/ S91C** | $0.63 \pm 0.02$ | $0.05 \pm 0.03$ | $0.44 \pm 0.02$ | $0.25 \pm 0.10$ | $0.26 \pm 0.02$ | $0.66 \pm 0.17$ |
| **C73/ C121** | $0.58 \pm 0.02$ | $0.14 \pm 0.08$ | $0.44 \pm 0.02$ | $0.34 \pm 0.10$ | $0.51 \pm 0.02$ | $0.27 \pm 0.08$ |

**Figure 5.**
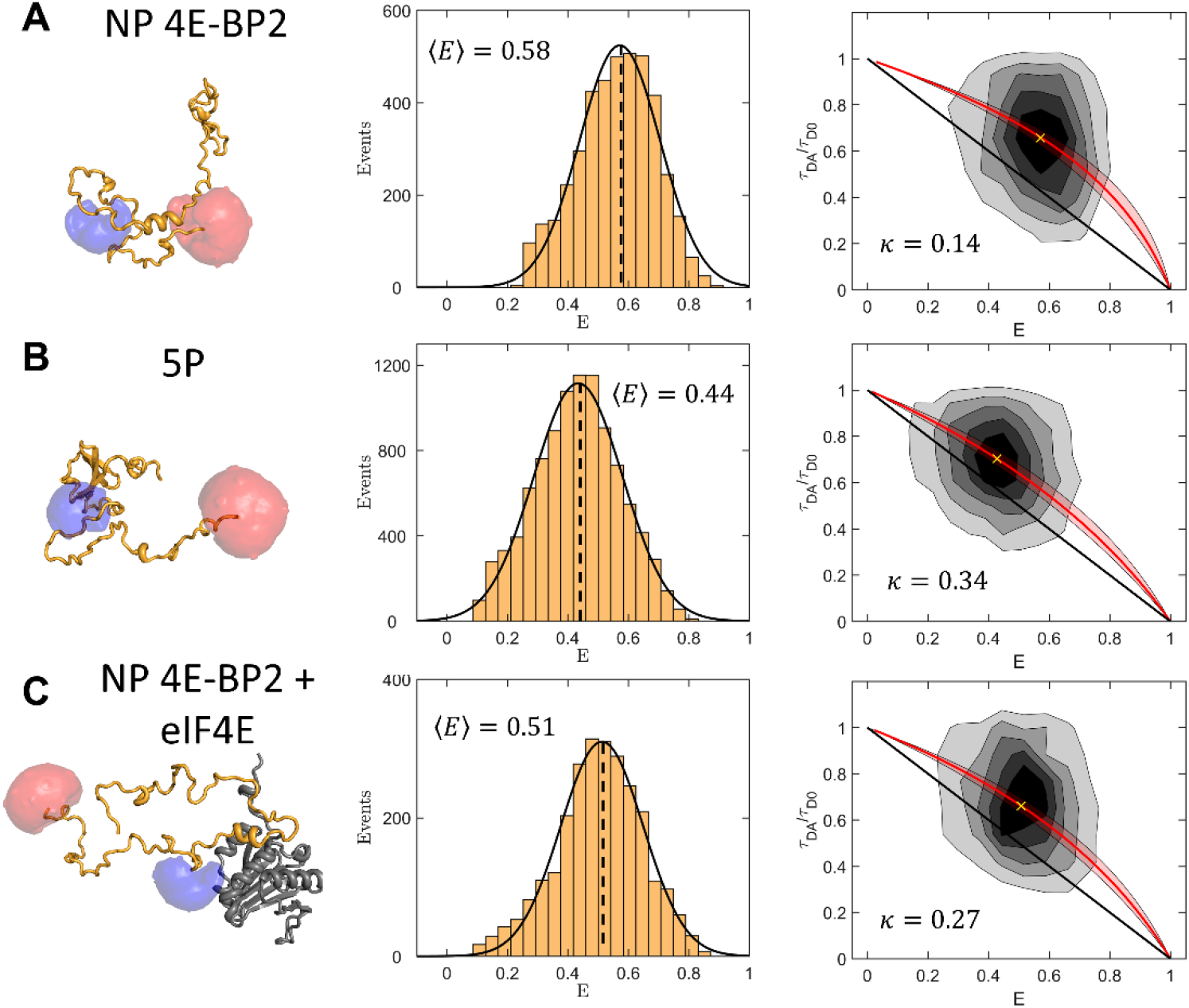
smFRET MPF results for the C73/C121 4E-BP2 construct labelled with Alexa Fluor 488 and Alexa Fluor 647 in the (**A**) non-phospho (NP), (**B**) five-phospho (5P) and (**C**) non-phospho eIF4E bound states (NP 4E-BP2 + eIF4E). (1^st^ column) Cartoon depictions of conformations of 4E-BP2 in different states with representations of fluorophore accessible volume simulations (47). (2^nd^ column) FRET efficiency histograms fit to a Gaussian function, with the dashed lines indicating the mean values. (3^rd^ column) 2D histograms plots of the FRET efficiency (*E*) and the donor-only normalized fluorescence lifetime; τ_DA_ and τ_D0_ are donor lifetimes in the presence and absence of acceptor, respectively. The black line shows the relation expected for a static rigid molecule. The red line is a relation for a worm-like chain with a stiffness parameter (κ) that passes through the centroid of a 2D gaussian fit indicated by a yellow cross. The red shaded region shows the uncertainty range of κ.

The FRET efficiency of the NP H32C/S91C construct exhibits an even larger decrease upon binding to eIF4E, i.e., 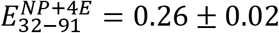. This construct flanks both primary and secondary binding sites and should be sensitive to 4E-BP2:eIF4E interactions and the 3D arrangement of the complex. The large FRET decrease could result from a combination of more extended conformations being compatible with the eIF4E bound state, in which 4E-BP2 wraps around eIF4E (with the exception of the canonical helical element), and an excluded-volume effect exerted by eIF4E on 4E-BP2. The conformations of 4E-BP2 bound to eIF4E are thought to be defined by dynamic interactions of the canonical and secondary binding elements, and by significant structural disorder elsewhere (20).

smFRET histograms of the C73/C121 construct for the same three 4E-BP2 states are shown in **Fig. 5**. The average FRET efficiency for NP 4E-BP2, 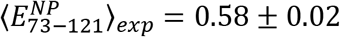, is also higher than that expected from a statistical coil, 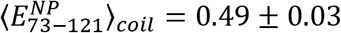, suggesting that the C-terminal disordered region contains transient intrachain contacts. The average C73/C121 FRET efficiency decreases to 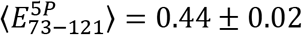 upon phosphorylation, but only to 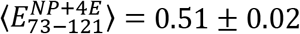 upon binding to eIF4E. The electrostatic repulsion between the five phosphates may contribute to the C-IDR expansion of 5P 4E-BP2. This expansion of the C-IDR has also been observed from previously published NMR paramagnetic relaxation enhancement (PRE) data and chemical shift-derived measures of secondary structure. (23)

For the bound state, the reduction in FRET is much less for C73/C121 than for H32C/S91C, i.e., 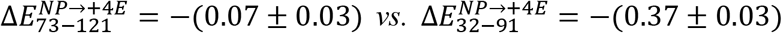. This suggests that C-terminal disordered conformations are less restricted in the NP 4E-BP2:eIF4E complex than those of the 32-91 region, as expected from the complete loss or very significant broadening of NMR resonances upon binding from residues 34-90 (20).

The relation between the donor lifetime and the FRET efficiency can be used to infer information about the dynamic exchange of the underlying states (50). A linear relation is expected for a static structure, with conformations that are rigid or fluctuate on a timescale slower than ~100 μs; in contrast, a nonlinear relation is expected for IDPs, as the burst duration (~1 ms) is much longer than typical chain reconfiguration times (~100 ns) (51). A family of dynamic *τ vs. E* lines based on a worm-like chain (WLC) model with variable stiffness *κ* (or persistence length) was generated using a method described by Barth et al. (52) The center of the experimental 2D FRET histogram was best matched to a WLC curve to infer the average stiffness for different regions of 4E-BP2 in different phospho/binding states (**Table 3**).

The stiffness parameter of the 32-91 region, 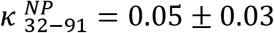, corresponds to a persistence length of 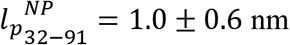, which is similar to *l_p_* = 0.4 ± 0.07 nm reported for a set of disordered and unfolded proteins.(53) The stiffness increases to 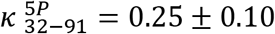 upon phosphorylation, and even more significantly, to 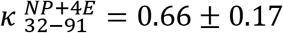 upon binding to eIF4E (**Fig. 4** and **Table 3**). The larger stiffness in the 5P state is consistent with the appearance of a stable beta-fold between residues 18 and 62. In the bound state, the 2D FRET population lies close to the static line; this is consistent with the decreased dynamics expected as residues 49-67 form a predominantly stable helix when 4E-BP2 is bound to eIF4E.(20, 54) At the same time, the separation between the static and dynamic regimes is much reduced in the low *E* range, which increases the uncertainty in estimating the chain stiffness.

The 73-121 region has a stiffness of 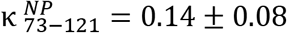 and 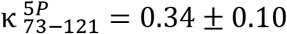 in the NP and 5P states, respectively. While slightly larger, these values mirror the changes observed for the 32-91 region. The stiffening of the C-IDR upon phosphorylation can be attributed to stabilizing interactions with the folded domain established previously (23). However, in contrast to the 32-91 region, the stiffness of the C-terminal region shows only a moderate increase in the eIF4E-bound state, to 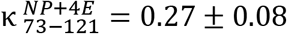, as the 2D FRET histogram remains well separated from the static line (**Fig. 5**). Together with the FAD and PET-FCS results, this suggests that the 73-121 region remains disordered and interacts with eIF4E as part of a more extensive dynamic complex than the current bipartite model (20).

## CONCLUSIONS

Dynamics is likely the key factor in understanding how 4E-BP2 regulates cap-dependent translation via interactions with the initiation factor eIF4E. Complementary to our previous NMR studies of the 4E-BP2 protein (12, 20, 23), a suite of multiparameter fluorescence techniques was used here to define local and global conformational dynamics of this intriguing IDP in its various states. Polarization anisotropy (FAD) measurements showed evidence of heterogeneous chain flexibility within the 4E-BP2 sequence, which correlates with variations in transient secondary structure and local amino acid composition. Multisite phosphorylation decreased segmental flexibility only in the region that undergoes folding (residues 18-62), while it made the rest of the chain more flexible. As anticipated, in the presence of eIF4E, segmental dynamics near the two binding sites was slowed down by binding interactions. Surprisingly however, the C-terminal region adjacent to the secondary binding site experienced the largest changes, while the N-terminal region exhibited much less change. These results implicate a much larger region of 4E-BP2 interacting with eIF4E than previously thought and showcase FAD as a sensitive method to probe local binding affinity, at the level of individual protein segments, in the context of multivalent interactions.

As seen previously for other IDPs (42), the quenching rates decreased at most labelling sites. This likely arises from a combination of local steric and electrostatic effects of the phosphate groups and the overall steric effect of the folded domain. In the presence of eIF4E, the quenching rates near the two binding motifs in 4E-BP2 increased significantly, as expected from increased contacts with several exposed tryptophan and tyrosine residues along the extensive binding interface of eIF4E. Although contributions from intra- and inter-molecular contact could not be distinguished, the significant changes observed at the C-terminus support the involvement of the C-terminal region of 4E-BP2 in dynamic interactions with eIF4E.

smFRET measurements informed on changes in 4E-BP2 chain dimensions and stiffness of central and C-terminal regions of the protein following phosphorylation or binding. In the NP state, as a consequence of transient intrachain secondary and tertiary contacts, the donor-acceptor distance turned out smaller (higher FRET efficiency) than the random coil prediction for both FRET constructs. In the 5P state, the separation between residues 32 and 91, which partially contains the folding domain, increased, instead of the decrease typically expected for a folding transition. When bound to eIF4E, the chain expands and stiffens considerably around the canonical and secondary binding sites, while the C-terminal region remains highly flexible. The canonical binding motif and secondary binding site provide specificity, while the dynamic nature of the complex which retains significant chain flexibility minimizes the entropic penalty. All phosphorylation sites except S65, which is phosphorylated last, are in regions that remain highly dynamic in the 4E-BP2:eIF4E complex. This facilitates the access of large kinases and allows a rapid response to cellular conditions. Our fluorescence-based characterization of the 4E-BP2 conformational dynamics is an important step towards understanding the interplay between folding and release of binding to eIF4E and its regulatory function, and it provides a foundation for future studies of IDP conformational and binding equilibria.

## MATERIALS AND METHODS

### Materials

The fluorescent probes used for labelling the 4E-BP2 protein were: Alexa Fluor 488 (A488) maleimide, Alexa Fluor 647 (A647) maleimide (ThermoFisher Scientific, Canada) and, Atto488 (At488) maleimide (ATTO-TEC GmbH, Germany). Guanidinium chloride (GdmCl) (G9284, Sigma Aldrich) was used for protein denaturation. All samples were diluted in phosphate-buffered saline (PBS) containing 137 mM NaCl, 2.7 mM KCl, 10 mM Na_2_HPO_4_, and 1.8 mM KH_2_PO_4_ at pH 7.4. GdmCl solutions were adjusted to pH 7.4 for all the denaturation experiments.

### Protein expression and purification

Small ubiquitin-like modifier (SUMO) fusion constructs of both the 4E-BP2 protein (residues 1 through 120) and eIF4E protein (residues 1 through 217) were expressed and purified as described previously(23). Briefly, the proteins were expressed in BL21-codonplus (DE3)-RIPL competent *E. coli* cells (Agilent Technologies) in Lysogeny broth at 37 °C until OD_600_~0.6-0.8, induced with isopropyl β-D-1-thiogalactopyranoside (IPTG), and expressed at 16 °C for ~16 h. Protein was purified from cell lysate with a nickel–nitrilotriacetic acid (Ni–NTA) column followed by cleavage of the SUMO solubility tag with ULP1 at 4 °C for ~16 h. The Sumo tag was separated using an Ni-NTA column followed by HiLoad Superdex 75 PG gel filtration column (28-9893-34, GE Healthcare) if the protein was not pure as assessed by SDS-PAGE. The molecular mass and the purity of protein samples were verified by electrospray ionization mass spectrometry (ESI-MS).

Phosphorylation of 4E-BP2 with activated Erk2 using a dialysis technique was performed as described previously(12, 23). Briefly, 50 mL of 5 μM Erk2 and 20 μM of 4E-BP2 were dialyzed in a 3 kDa MWCO dialysis bag in 1 L of buffer. The dialysis buffer contained 50 mM Tris-HCl, pH 7.5, 1 mM EGTA, 5 mM BME, 20 mM MgCl_2_, 10 mM EDTA, and 10 mM ATP, dialysis was performed at 20 °C for 1-3 days. Phosphorylated 4E-BP2 was purified from Erk2 using a Ni-NTA column. Purity and degree of phosphorylation of 4E-BP2 was confirmed by ESI-MS.

All single cysteine proteins (C0/C35S/C73S (cysteine insertion at 0 position), S14C/C35S/C73S, C35/C73S, C73/C35S, C35S/C73S/S91C, C35S/C73S/C121 (cysteine insertion at 121 position)) were labeled by adding the Atto488 maleimide fluorophore to a 50 μL solution of 100 μM protein at a dye:protein molar ratio of 3:1. The double-cysteine mutants (H32C/C35S/C73S/S91C) and (C35S/insC121) was labeled with Alexa Fluor 488 maleimide and Alexa Fluor 647 maleimide by adding Alexa Fluor 488 and Alexa Fluor 647 to a 50 μL solution of 100 μM protein at a A488:A647:protein molar ratio of 1.3:3:1. TCEP was added at a 10× molar excess to the protein in order to reduce the disulfide bonds. All the maleimide-cysteine coupling reactions were performed in a PBS buffer at pH 7.4. Oxygen was removed by flushing the sample with argon gas in a desiccator for 5 min. The vial was capped tightly and shaken gently for 3 hours at room temperature. The excess dye was removed by size-exclusion chromatography using Sephadex G-50 gel (G5080, Sigma Aldrich) in a BioLogic LP system (731-8300, Bio-Rad).

All samples were diluted to concentrations of 1–10 nM and 20–50 pM, which are most suitable for FCS/FAD and smFRET burst experiments, respectively. For a typical experiment, a sample solution of 30 μL was dropped on the surface of plasma-cleaned coverslip. Non-specific protein adsorption to the coverslip was prevented by adding 0.005% (v/v) Tween-20 (P2287, Sigma-Aldrich) to the solution, and bovine serum albumin (BSA) (15260-037, ThermoFisher Scientific) was used to coat the clean coverslips. All experiments were performed at 20 °C.

### Instrumentation

smFRET measurements were performed on a custom-built multiparameter fluorescence microscope (32). The donor was excited at 480 nm by frequency doubling the infrared output of a femtosecond laser (Tsunami HP, Spectra Physics), while the acceptor was excited at 635-nm using a diode laser (WSTech, TECRL-25GC-635). Alternating-laser excitation (ALEX) of the sample was performed by synchronous modulation of the two laser sources to achieve alternating 50-μs periods of donor and acceptor fluorophore excitation, respectively. Laser intensities of 10 kW/cm^2^ and 3.6 kW/cm^2^ at the sample were used for exciting the donor and the acceptor fluorophores, respectively. FAD measurements were performed on the same microscope, by exciting Atto488 at an average intensity of ~0.14 kW/cm^2^ at the sample. FCS measurements were performed on a separate custom-built fluorescence microscope described elsewhere (55), where Atto488 was excited using a 488-nm diode laser (TECBL-488nm, WorldStarTech) at an average intensity of ~5 kW/cm^2^ at the sample.

### FAD analysis

FAD monitors the rotation dynamics of the emission dipole of the dye. The “wobble-in-a-cone” model (30) was used to fit the experimental FAD data:

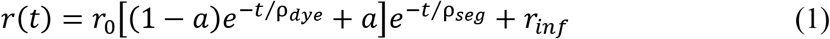

where *ρ_dye_*, *ρ_seg_* are rotational correlation lifetimes of the dye and the protein segment, respectively, *a* is the fraction of the slower (segmental) component, *r_0_* is initial (limiting) anisotropy, and *r_inf_* is the non-decaying (residual) anisotropy. The uncertainties of the fitted parameters were estimated by taking the standard deviation of the fitting results of 3-7 data sets collected consecutively; if any of the fitting derived errors were larger, then this value was reported. The baseline offset *r_inf_* accounts for the slow, global motion of the whole protein; this is typically very small for IDPs due to their high backbone flexibility (29), but it can increase upon binding to their targets.

### FCS analysis

In FCS, the fluorescence autocorrelation function for free Brownian diffusion of a single molecular species with multiple relaxation components is given by (57):

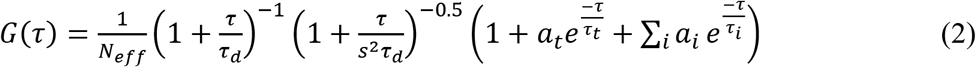

In equation (3), *N_eff_* is the average number of molecules in the detection volume, *s* is the ratio between the axial and the lateral radii of the detection volume (*s* = *z*_0_/*w*_0_), and *τ_d_* is the average diffusion time, which is related to the diffusion coefficient 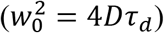 and to the hydrodynamic radius *R_H_* of the molecule via the Stokes-Einstein equation(58); *a_t_* and *τ_t_* are the amplitude and lifetime of the triplet component and *a_i_* and *τ_i_* are the amplitude and the lifetime of the *i*^th^ PET relaxation components, respectively. Global fitting was performed simultaneously on each labelling position (C0, C14, C35, C73, C91, C121), a separate global fit was performed for each for each measurement condition (NP 4E-BP2, 5P 4E-BP2 and NP 4E-BP2 + eIF4E), a total of three independent global fits performed on six FCS curves simultaneously. In the global analysis the diffusion time (*τ_d_*), the triplet amplitude (*a_t_*) and the triplet lifetime (*τ_t_*) were shared parameters, while the average number of molecules in the detection volume (*N_eff_*), the PET relaxation amplitudes (*a_i_*) and lifetimes (*τ_i_*) were fit as individual parameters. PET relaxation components with amplitudes of 0.05 or less were considered insignificant and not reported. The triplet lifetime (*τ_t_*) were confined to a range of 5-10 μs based on characterization of the triplet state of free Atto488 as a function of excitation intensity, see **Fig. S5**. FCS measurements performed at ~5 kW/cm^2^, returned a characteristic diffusion time (*τ_d_*) for free Atto488 of 48 ± 1 μs. To exclude the possibility that the slow relaxation component (*τ*_1_) resulted from the diffusion of unreacted free Atto488 remaining after size-exclusion chromatography purification, we performed an additional reverse-phase high-performance liquid chromatography (RP-HPLC) purification of the S91C NP 4E-BP2 construct. We did not detect unreacted Atto488 in the RP-HPLC chromatogram (**Fig. S7**), we also observed the slow relaxation component (*τ*_1_) in FCS measurements of the S91C NP 4E-BP2 after RP-HPLC purification (**Fig. S8**). Fitting was optimized by varying the number of relaxation components, the goodness of fit (*χ*^2^) and Akaike information criterion (AIC) were calculated for each fit. The addition of relaxation components was accepted if: the *χ*^2^ parameter decreased, the AIC decreased, and the fitting residuals were featureless. The uncertainties of the fitted parameters were estimated using the Jacobian from Levenberg-Marquardt least-squares fitting. Prior to each set of measurements, a calibration dye (rhodamine 110) was used to estimate the *s* and w_0_ parameters, typically ~8 and ~250 nm, respectively.

### smFRET analysis

A custom MATLAB script based on the ‘MLT’ algorithm was used to identify fluorescence bursts and sort them into donor-only, acceptor-only and dual-labelled (FRET) populations (32). The FRET efficiency for each burst was calculated based on the number of detected photons in the donor (*I_D_*) and acceptor (*I_A_*) channels:

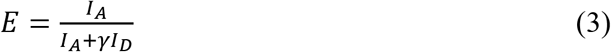

where *γ* is a correction factor for the difference in the detection efficiencies of donor and acceptor channels, and the quantum yields of the dyes were determined as described in SI section 2.1. In addition, corrections were applied on both *I_D_* and *I_A_* to subtract background, spectral cross talk, and direct excitation of the acceptor. The background was obtained from a measurement of the sample buffer while the corrections for cross talk and direct excitation were derived from donor-only and acceptor-only bursts. A smFRET histogram was constructed from all bursts detected for a given sample, each histogram was fit to a single Gaussian distribution.

To estimate the FRET efficiency if 4E-BP2 was a featureless statistical coil, 5000 conformers were generated using Trajectory Directed Ensemble Sampling (TraDES)(59) with 100% coil sampling in accordance with the sequences in **Table S1**. The FRET efficiency for each conformer was calculated using the python library *LabelLib*(60) (see SI section **2.3**).

For each FRET construct, a family of τ_DA_/τ *vs E* curves were generated based on a WLC model with stiffness parameters *κ* between 0.01 and 1 using the *FRETlines* Python library(52). The chain stiffness for given dataset was selected as the *κ* value of the τ_DA_/τ *vs E* curve passing through the centroid of a 2D Gaussian fit of the experimental τ_DA_/τ *vs E* histogram. The confidence interval was established from the range of WLC models that agree with the centroid from fitting within the experimental accuracy of determining the FRET efficiency, i.e., 0.02 (32).

## AUTHOR INFORMATION

### Author Contributions

C.C.G. was responsible for overall project management and supervision. S.S. and Z.Z. expressed, purified and labelled proteins, performed measurements and analyzed data. A.B. and J.D. designed single- and double-cysteine 4E-BP2 mutants. T.T. performed smFRET simulations. S.S., Z.Z., and C.C.G. wrote the manuscript. J.D.F.-K. contributed to interpretation of results and critical revision of manuscript.

## ACKNOWLEDGEMENTS

This work has been supported by the Natural Sciences and Engineering Research Council of Canada (NSERC RGPIN 2017 – 06030 to C.C.G.), the Canadian Institutes of Health Research (CIHR FND-148375 to J.D.F.-K.) and National Institutes of Health (NIH 5R01GM127627-03 to J.D.F.-K.).

