## Supplemental Information for "Multisite Phosphorylation and Binding Alter Conformational Dynamics of the 4E-BP2 Protein"

### SUPPLEMENTAL INFORMATION SECTION 1:
### Expression, Purification, Labelling, and Phosphorylation of 4E-BP2 and eIF4E

#### 1.1 4E-BP2 Expression, Purification and Labelling

A single bacterial colony on kanamycin LB Agar plates was inoculated into 5 mL of LB containing 50 mg/L kanamycin and cultured at 37 ºC overnight. This medium was used to inoculate 1 L of kanamycin containing LB and grown at 37 ºC until $OD_{600}$~0.6-0.8. Cells were induced with IPTG to a concentration of 1 mM and grown for ~16 h at 16 ºC.

Cells pellets were harvested by centrifugation at (3214×g, 30 minutes, 4 ºC), resuspended in a lysis buffer (20 mM $NaH_2PO_4$, 500 mM NaCl, 5 mM BME, 20 mM Imidazole, 6 M GdmCl, pH 7.5) and lysed by sonication. Soluble components were purified from insoluble by centrifugation (30000×g, 60 minutes, 4 ºC), the soluble fraction was bound to a nickel–nitrilotriacetic acid (Ni–NTA) column for 30 minutes. A thorough washing with the lysis buffer to remove non-specifically bound proteins was followed by elution with the elution buffer (20 mM $NaH_2PO_4$, 500 mM NaCl, 5 mM BME, 400 mM Imidazole, pH 7.5). The elution fraction was placed in a dialysis bag (3 kDA MWCO) with ~100 µg ULP1 at 4 ºC for ~16 h in 4L of dialysis buffer (50 mM Tris-HCl, 200 mM NaCl, 5 mM BME, 10 mM Imidazole, pH 7.5) to cleave the His-SUMO tag. GdmCl was added to the cleavage products to a concentration of 2 M and incubated on a Ni-NTA column for 30 minutes. 4E-BP2 was purified from ULP1 and the His-SUMO tag by washing with the dialysis buffer containing 2 M GdmCl, the purity was assessed by SDS-PAGE. If required further purification was performed with a HiLoad Superdex 75 PG gel filtration column in the dialysis buffer with 2 M GdmCl.

Immediately prior to the labelling, the protein was buffer exchanged and concentrated at 4 ºC (Amicon Ultra-0.5 mL, UFC500396) to 100 µM in degassed labeling buffer (137 mM NaCl,

2.7 mM KCl, 10 mM $Na_2HPO_4$, and 1.8 mM $KH_2PO_4$, 1 mM TCEP, pH 7.4). Single-cysteine mutants were labelled with Atto488 maleimide (ATTO-TEC GmbH, Germany) at a dye:protein molar ratio of 3:1, double-cysteine mutants were labelled with Alexa Fluor 488 C5 Maleimide (ThermoFisher Scientific, Invitrogen, A10254) and Alexa Fluor 647 C2 Maleimide (ThermoFisher Scientific, Invitrogen, A20347) at a Alexa 488:Alexa 647:protein molar ratio of 1.3:3:1. Oxygen was removed by flushing the sample with argon gas in a desiccator for 5 min. The vial was capped tightly and shaken gently for 3 hours at room temperature. The excess dye was removed by size-exclusion chromatography using Sephadex G-50 gel (G5080, Sigma Aldrich) in a BioLogic LP system (731-8300, Bio-Rad). For dually labeled samples, the unlabeled and labeled species were further separated by MonoQ 5/50 GL column (GE Healthcare) in the AKTA protein purification system (18190026, GE Healthcare). Labelled samples were measured within 24 hours or stored immediately at -80 ºC for later use.

#### 1.2 4E-BP2 Phosphorylation

Phosphorylation of 4E-BP2 with activated Erk2 using a dialysis technique was performed as described previously(1, 2). Briefly, 50 mL of 5 µM Erk2 and 20 µM of 4E-BP2 were dialyzed in a 3 kDa MWCO dialysis bag in 1 L of buffer. The dialysis buffer contained 50 mM Tris-HCl, pH 7.5, 1 mM EGTA, 5 mM BME, 20 mM $MgCl_2$, and 10 mM EDTA, dialysis was performed at 20 ºC for 1-3 days. Phosphorylated 4E-BP2 was purified from Erk2 using a Ni-NTA column. Purity and degree of phosphorylation of 4E-BP2 was confirmed by ESI-MS. Phosphorylated samples were measured within 24 hours or stored immediately at -80 ºC for later use.

#### 1.3 eIF4E Expression and Purification

A single bacterial colony on kanamycin LB Agar plates was inoculated into 5 mL of LB containing 50 mg/L kanamycin and cultured at 37 ºC overnight. This medium was used to inoculate

1 L of kanamycin containing LB and grown at 37 ºC until $OD_{600}$~0.6-0.8. Cells were induced with IPTG to a concentration of 1 mM and grown for ~16 h at 16 ºC.

Cells pellets were harvested by centrifugation at (3214×g, 30 minutes, 4 ºC), resuspended in a lysis buffer (20 mM $NaH_2PO_4$, 500 mM NaCl, 5 mM BME, 20 mM Imidazole, 1 mM EDTA, DNASE1, protease Inhibitor cocktail cOmplete (Roche Diagnostics, Laval, Quebec, Canada), pH 7.5) and lysed by sonication. Soluble components were purified from insoluble by centrifugation (30000×g, 60 minutes, 4 ºC), the soluble fraction was bound to a Ni–NTA column for 30 minutes. A thorough washing with the lysis buffer to remove non-specifically bound proteins was followed by elution with the elution buffer (20 mM $NaH_2PO_4$, 500 mM NaCl, 5 mM BME, 400 mM Imidazole, pH 7.5). The elution fraction was placed in a dialysis bag (3 kDA MWCO) with ~100 µg ULP1 at 4 ºC for ~16 h in 4L of dialysis buffer (50 mM Tris-HCl, 200 mM NaCl, 5 mM BME, 10 mM Imidazole, pH 7.5) to cleave the His-SUMO tag. Cleavage products were incubated on an Ni-NTA column for 30 minutes. 4E-BP2 was purified from ULP1 and the His-SUMO tag by washing with the dialysis buffer, further purification was performed with a HiLoad Superdex 75 PG gel filtration column in a buffer containing 50 mM $NaH_2PO_4$, 150 mM NaCl, 5 mM BME, 2 mM Benzamidine-HCl, 1 mM EDTA, at pH 7.5.

### SUPPLEMENTAL INFORMATION SECTION 2:

### Single-Molecule Förster Resonance Energy (smFRET)

#### 2.1 Instrumental Gamma Correction Factor: Double-stranded DNA

The gamma correction factor accounts for the detection efficiency of the instrument and quantum yield of the dyes. We designed three 30 base-pair (bp) double-stranded DNA (dsDNA) constructs with expected FRET efficiencies of ~30%, 50%, and ~80%, with bp separations of 17, 13, and 10 respectively. Complementary single-stranded DNA (ssDNA) strands with the donor (fluorescein) conjugated to a thymine ring on one strand and the acceptor (Alexa Fluor 647) conjugated to the 5'-end of the complementary strand were synthesized by Integrated DNA Technologies (IDT). Equimolar amounts (100 μM) of the two strands were heated to 95 ºC for 2 minutes using a thermocycler (Biometra TPersonal Thermocycler) in the annealing buffer (10 mM Tris, 50 mM NaCl, 1 mM EDTA, pH 7.5), followed by slow cooling to 25 ºC for 45 minutes, after which the sample was aliquoted and kept at -20 ºC.

The gamma factor for the dsDNA was determined using the three constructs by fitting the relationship between $\langle E_{PR} \rangle$ and $\langle S_{PR} \rangle^{-1}$ as described by Kapanidis et. al.(3). The DNA gamma factor was used to estimate the sample gamma factor: $\gamma_{sample} = \gamma_{DNA} \frac{\phi_A^{sample} \phi_D^{DNA}}{\phi_D^{sample} \phi_A^{DNA}}$, where the quantum yields of the donor and acceptor of all samples were estimated using the lifetime method, $\frac{\tau_x}{\tau_y} = \frac{\phi_x}{\Phi_y}$.

### 2.2 Förster Radius Estimation

Donor emission and acceptor absorption spectra of the H32C/S91C 4E-BP2 FRET construct, were measured on donor-only and acceptor-only labelled proteins, labelled with Alexa Fluor 488 (A488) and Alexa Fluor 647 (A4647) respectively, which were prepared using the protocol described in **SI section 1.1**. The Förster radius for this construct, $R_0$, was estimated as:

$$R_0{}^6 = \frac{9000 \cdot ln(10) \cdot \Phi_D \kappa^2}{128 \pi^5 N_A n^4} \int_0^\infty f_D(\lambda) \varepsilon_A(\lambda) \lambda^4 d\lambda \quad \text{(S1)}$$

Where $\Phi_D$ is the donor fluorescence quantum yield, $\kappa^2$ is the dipole orientation factor, $N_A$ is Avogadro's number, $n$ is the refractive index, and $J$ is the normalized spectral overlap integral $J =$

$\int_0^\infty f_D(\lambda)\varepsilon_A(\lambda)\lambda^4 d\lambda$. The quantum yield of the donor was determined by the lifetime method (**SI section 2.1**), the isotropic orientation factor was used ($\kappa^2$ = 2/3), and the refractive index was assumed to be that of water ($n = 1.33$). The $\kappa^2$ = 2/3 assumption is supported by the anisotropy decay features: fast depolarization relative to the fluorescence lifetime and very small values of the residual anisotropy (**Table S2**). The Förster radius was estimated to be 51 ± 2 Å for the H32C/S91C 4E-BP2 in the NP state, labelled stochastically with A488 and A647. For the equivalent C73/C121 FRET construct, the spectral overlap integral was assumed to be the same as that of H32C/S91C. In addition, there was no change in quantum yield of the donor, $\Phi_D$, for H32C/S91C and C73/C121 in equivalent phosphorylation states, and therefore the same Förster radii were used for the latter FRET construct.

#### 2.3 Prediction of the 4E-BP2 Random Coil FRET Efficiency

Random coil (RC) conformations of the 4E-BP2 sequence (Fig. S1) were created with the statistical coil generator Trajectory Directed Ensemble Sampling (TraDES)(4), which imposes a Lennard-Jones type potential to avoid steric clashes between residues. The algorithm requires a trajectory file created using the program seq2trj. The trajectory file defines a probability distribution of peptide dihedral angles to be sampled, which are grouped into alpha, extended/beta-sheet and coil. The RC ensemble for NP 4E-BP2 included 5000 conformers which were generated using TraDES by sampling exclusively coil-type dihedral angles.

The FRET efficiency, $E$, for each conformation within the RC ensemble was estimated as:

$$E = \frac{1}{\left[1+\left(\frac{r}{R_0}\right)^6\right]} \tag{S2}$$

where $r$ is the donor-acceptor distance and $R_0$ is the Förster radius calculated above, 51 ± 2 Å (**section SI 2.2**). The donor-acceptor distance ($r$) was sampled by Accessible Volume (AV) simulations(5) of the two fluorophores with their linkers attached to designed labeling positions for each FRET construct. The geometrical dimensions of the two fluorophores, Alexa Fluor 488 and Alexa Fluor 647, in the AV simulations were as described previously(5). The uncertainty of the computed average FRET efficiency (±0.03) was obtained by propagating the uncertainty of $R_0$. The standard error of the mean calculated was ~0.003 for both FRET constructs, which is noticeably less than the error propagated from $R_0$. The computations were performed with AvTraj (5) v0.0.9 and MDTraj (6) v1.9.3 packages in Python 3.7.6.

#### 2.4 Burst Search, Filtering, and Corrections

Multiparameter fluorescence analysis (7, 8) and alternating-laser excitation (9) (ALEX) based filtering was applied to all acquired data. An all photon burst search(10, 11) with $M$ = 30, $L$ = 30, and $T$ = 500 µs was used to identify bursts from the photon stream. Where M is the minimum number of photons per averaging time window, L is the minimum number of photons per burst, and T is the averaging time window. Stoichiometry values, as defined in **Eq. S3**, were computed for each burst, where number of photons detected in donor and acceptor channels after donor excitation are $I_{D_{ex}}^{D_{em}}$ and $I_{D_{ex}}^{A_{em}}$, respectively, $I_{A_{ex}}^{A_{em}}$ is the number of photons in acceptor channel after acceptor excitation. Only bursts with stoichiometry between 0.1 and 0.7 ($0.1 < S < 0.7$) were analyzed.

$$S = \frac{I_{D_{ex}}^{D_{em}} + I_{D_{ex}}^{A_{em}}}{I_{D_{ex}}^{D_{em}} + I_{D_{ex}}^{A_{em}} + I_{A_{ex}}^{A_{em}}} \qquad \text{(S3)}$$

The background was obtained from a measurement of the sample buffer, background was subtracted from each detector, separate corrections were applied to each detection channel and whether donor or acceptor excitation was used. Leakage ($Lk$) and direct-excitation ($Dir$) correction factors were obtained from donor-only and acceptor only burst respectively, typical values were $Lk$~0.03 and $Dir$~0.05. The leakage and direct-excitation correction factors were applied to **Eq. 3**, main text, using $F_{Lk} = Lk \cdot n_D$ and $F_{Dir} = Dir \cdot n_A$ respectively. The gamma correction factor was obtained as described in SI **section 2.1** and applied to **Eq. 3**, main text.

#### 2.5 Two-dimensional Lifetime vs. FRET Efficiency Analysis

For each FRET construct, a family of $\tau_{DA}/\tau$ *vs E* curves were generated based on a WLC model with stiffness parameters $\kappa$ between 0.01 and 1 using the *FRETlines* Python library (12). The chain stiffness for given dataset was selected as the $\kappa$ value of the $\tau_{DA}/\tau$ *vs E* curve with the smallest difference $\sqrt{(\Delta E)^2 + \left(\Delta \frac{\tau_{DA}}{\tau}\right)^2}$ from the centroid of a 2D Gaussian fit of the experimental $\tau_{DA}/\tau$ *vs E* histogram. The confidence interval was established from the range of WLC models that agree with the centroid from fitting within experimental FRET efficiency uncertainty (**Fig. 4** and **Fig. 5**). Assuming an inter-residue distance of 3.4 Å, the contour length of the H32C/S91S segment was estimated to be $L$ = 20.1 nm. Given that $\kappa = \frac{l_p}{L}$, the persistence length associated with the estimated $\kappa_{32-91}^{NP} = 0.05 \pm 0.03$ is ${l_p}_{32-91}^{NP} = 1.0 \pm 0.6$ nm, where $l_p$ is the persistence length and $L$ is the contour length. Using the same method, the persistence length of the C73/C121 segment was estimated to be ${l_p}_{73-121}^{NP} = 2.3 \pm 1.3$ nm.

### SUPPLEMENTAL FIGURES

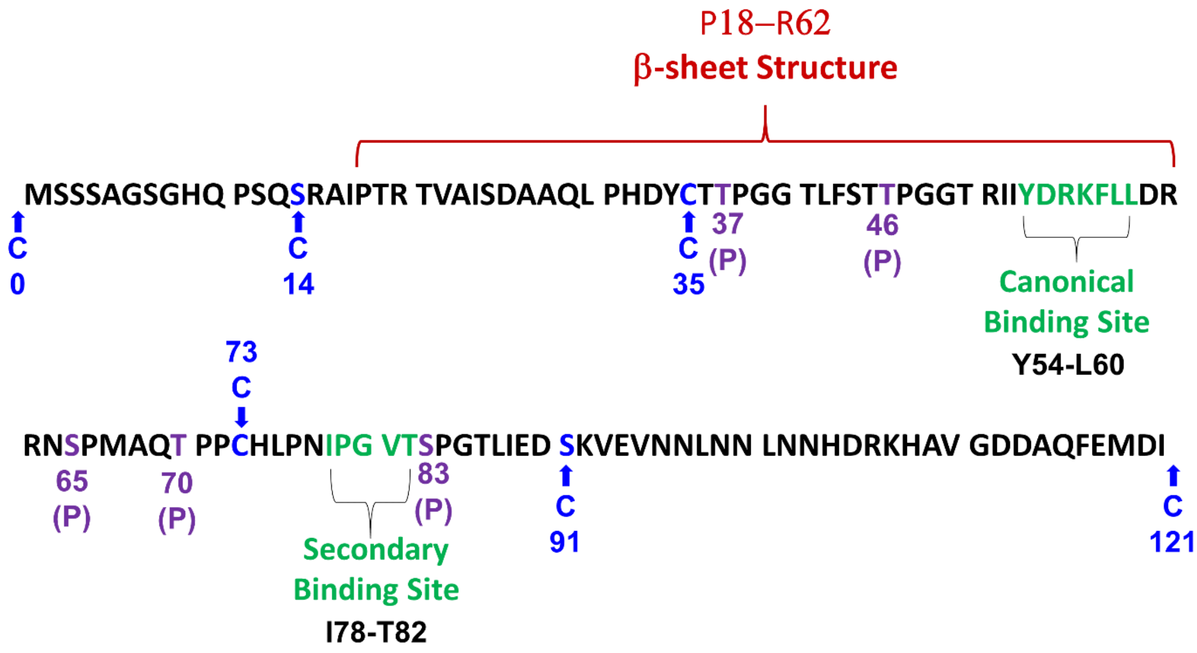


**Figure S1.** The sequence of human 4E-BP2 showing sites of cysteine maleimide fluorophore labelling in (**blue**) and sited of hierarchical phosphorylation in (**purple**). The canonical and secondary eIF4E binding sites are shown in (**green**), the residues involved in formation of the β-fold are outlined in (**red**).

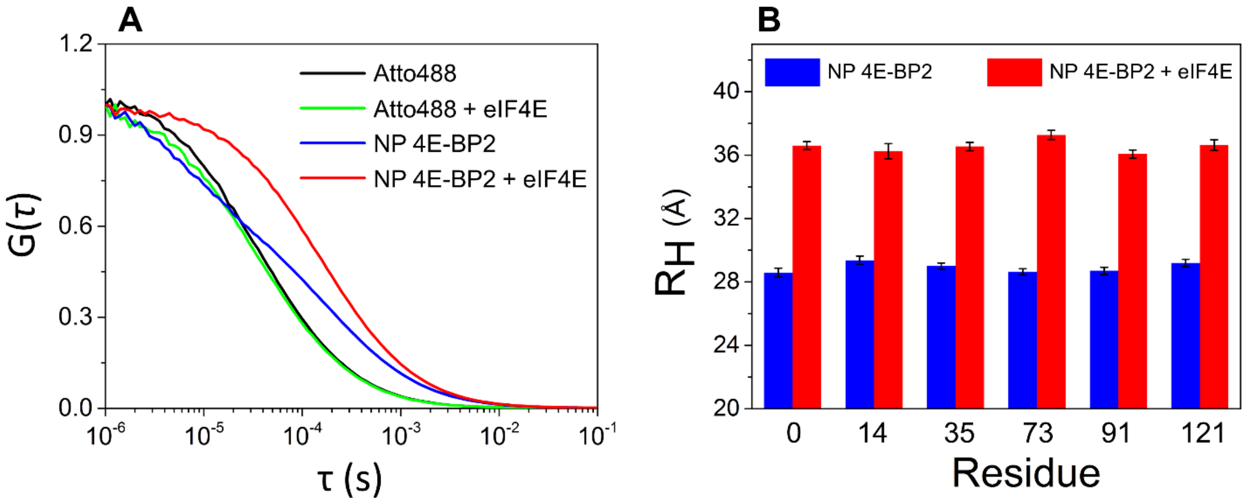


**Figure S2.** (**A**) Normalized experimental FCS curves for the At488 dye alone (**black**), a mixture of At488 and eIF4E (**green**), non-phosphorylated 4E-BP2 labeled with At488 at residue 73 in the absence (**blue**) and in the presence of 0.5 µM eIF4E (**red**). (**B**) Hydrodynamic radii of NP 4E-BP2 alone (**blue**) and eIF4E-bound (**red**), as estimated by fitting FCS data for different labelling positions to **Eq. 2**. Error bars are the standard deviations from fitting.

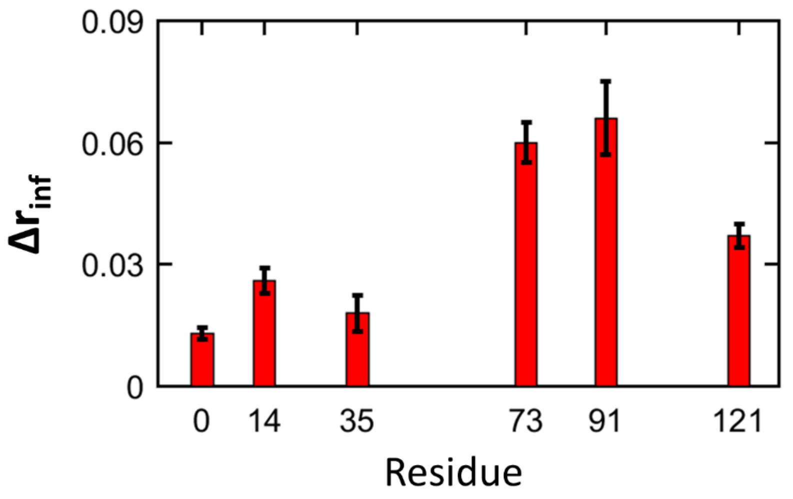


**Figure S3.** Changes in the residual anisotropy, $\Delta \boldsymbol{r_{inf}}$, upon binding of NP 4E-BP2 to eIF4E at different labelling sites in the 4E-BP2 sequence. Residual anisotropies were obtained from fitting the FAD decays at each labelling site to **Eq. 1** (see main text); results are listed in **Table S2**. Error bars are the propagated errors from standard deviations of the fitting.

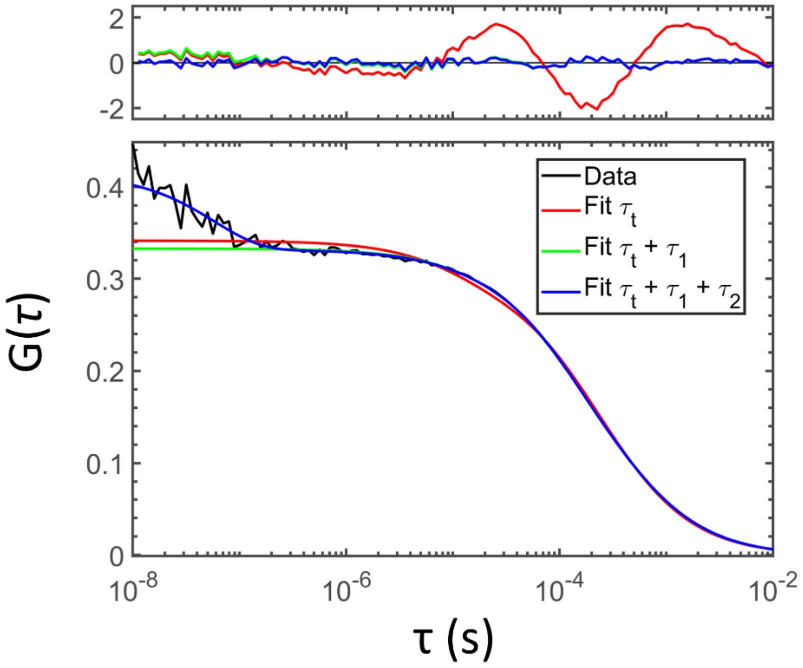


**Figure S4.** NP 4E-BP2 labeled at residue 73 in the presence of 0.5 µM eIF4E was fit to **Eq. 2** with only the triplet component, or with triplet component and an additional 1 or 2 kinetic decay components. Judging from the shape of the residuals and the direct comparison of the data and the fit, two kinetic components were required to satisfactorily fit the data in this case. The full results are given in **Table S3**.

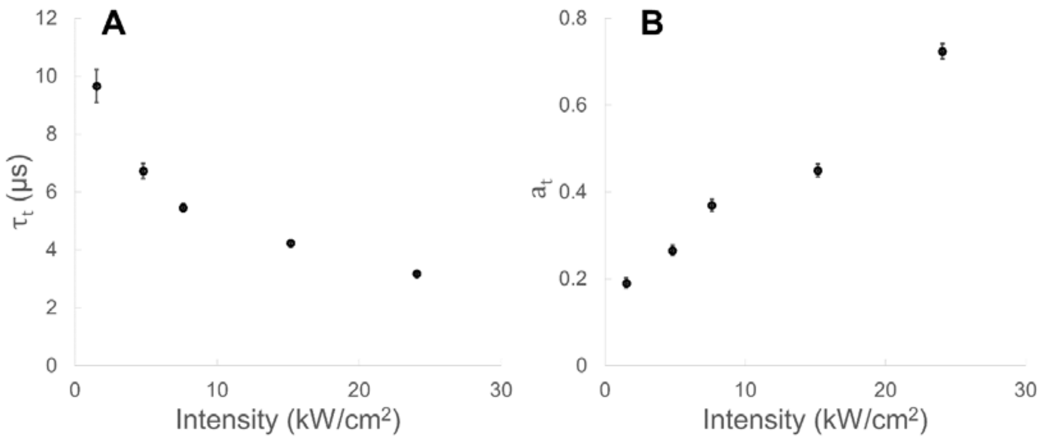


**Figure S5.** Triplet state parameters for Atto488 at different excitation intensities. The triplet lifetimes (**A**) and amplitudes (**B**) were obtained from fitting FCS curves to **Eq. 2** with one decay component ($\tau_t$) and one amplitude $a_t$. Error bars represent the standard deviation from fitting; some error bars are too small to be seen.

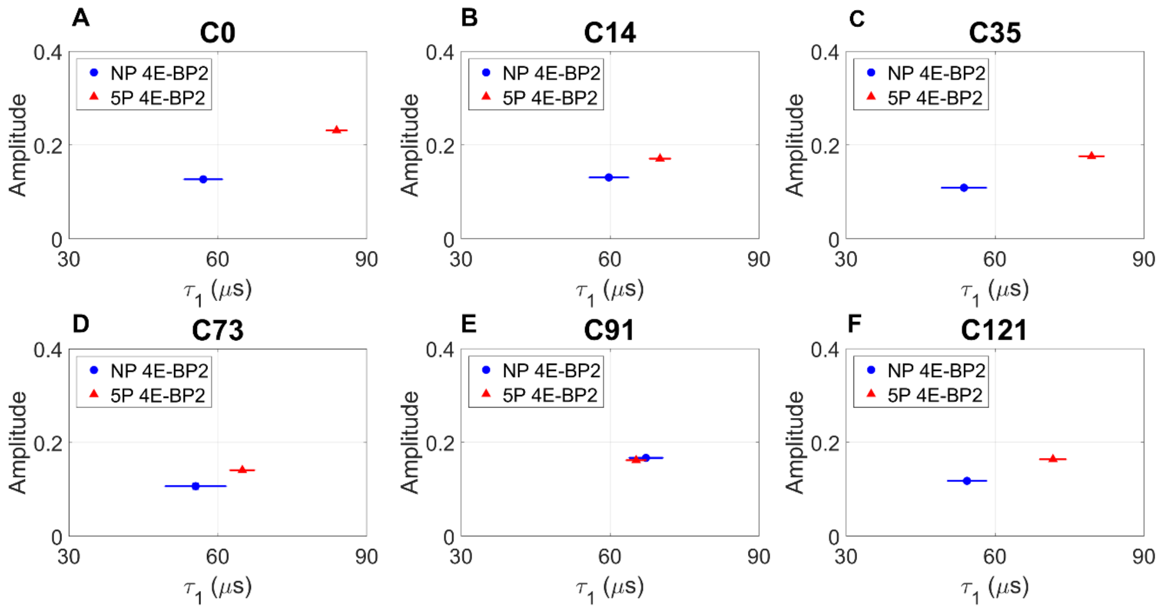


**Figure S6.** The changes in lifetime ($\tau_1$) and amplitude **(*$a_1$*)** of the slow FCS kinetic component upon multisite phosphorylation. 4E-BP2 was labelled with Atto488 at positions 0, 14, 35, 73, 91 and 121 along the sequence, and the experimental FCS curves were fit to **Eq. 2**. Different protein states are indicated as follows: non-phospho (**NP, blue circle**), five-phospho (**5P, red triangle**). The fitting error bars for each parameter are indicated in the figure; all fitting parameters are given in **Table S3**.

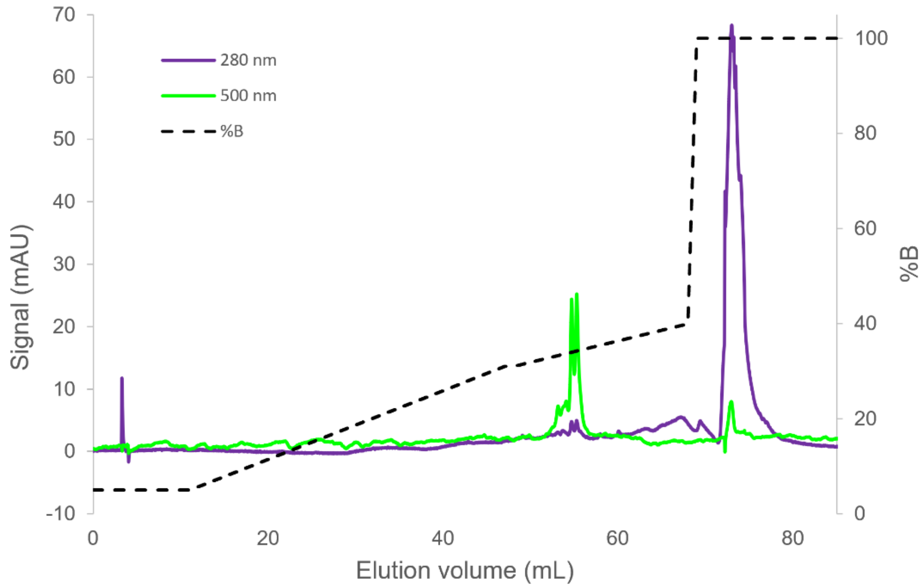


**Figure S7.** RP-HPLC purification of S91C NP 4E-BP2 labelled with Atto488, showing two major peaks, corresponding to the Atto488-labeled and unlabeled protein, which elute at ~55 mL and ~73 mL respectively. The labelled protein was initially purified from unreacted dye using size-exclusion chromatography with Sephadex G-50 gel. The small peak at which elutes at ~4 mL corresponds to sample which did not bind to the column during injection. Separations were performed on a C18 column at a flow rate of 1mL/min; solvent A is water + 0.1% TFA, and solvent B is acetonitrile.

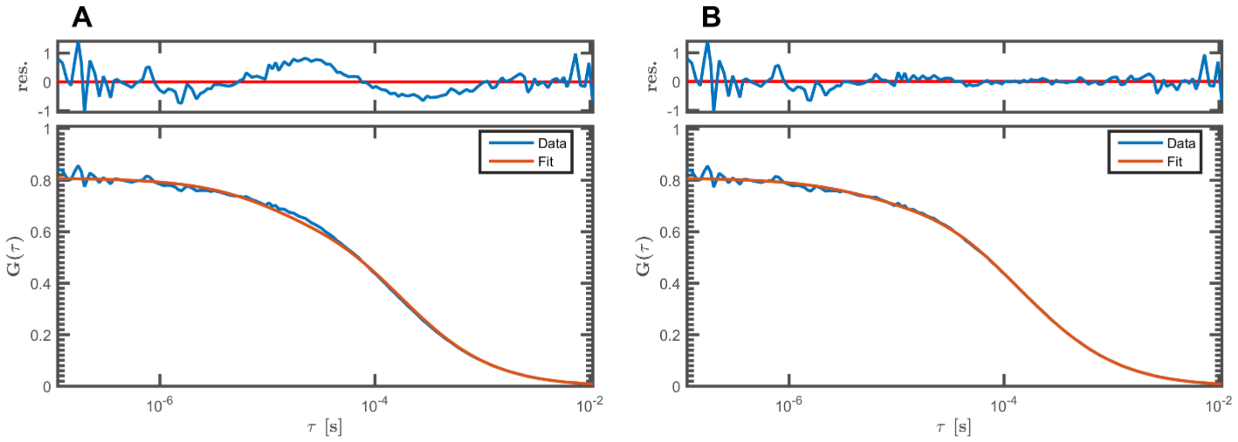


**Figure S8.** NP 4E-BP2 labelled with Atto488 at residue 91 after RP-HPLC purification (**Fig. S7**) fit to **Eq. 2** with **(A)** one decay component ($\tau_t$) accounting for the triplet state **(B)** and two decay components, one representing the triplet state ($\tau_t$) and one representing ($\tau_1$) the slow component.

### SUPPLEMENTAL TABLES

**Table S1.** Sequence mutations of 4E-BP2 and eIF4E.

| Protein | Construct | Mutations |
|---|---|---|
| 4E-BP2 | wt | - |
| 4E-BP2 | C0 | C0ins/C35S/C73S |
| 4E-BP2 | C14 | S14C/C35S/C73S |
| 4E-BP2 | C35 | C73S |
| 4E-BP2 | C73 | C35S |
| 4E-BP2 | C91 | C35S/ C73S/S91C |
| 4E-BP2 | C121 | C35S/ C73S/C121ins |
| 4E-BP2 | H32C/S91C | H32C/C35S/C73S/S91C |
| 4E-BP2 | C73/C121 | C35S/C121ins |
| eIF4E | I35C | I35C |

**Table S2.** Anisotropy decay parameters for different states of 4E-BP2 labelled at different sites along the sequence [a].

| | **C0** | **S14C** | **H32C** | **C35** | **C73** | **S91C** | **C121** |
|---|---|---|---|---|---|---|---|
| **NP** | | | | | | | |
| $\rho_{dye}$ | 0.51 ± 0.02 | 0.64 ± 0.03 | 0.35 ± 0.01 | 0.64 ± 0.05 | 0.62 ± 0.04 | 0.61 ± 0.04 | 0.53 ± 0.03 |
| $\rho_{seg}$ | 1.61 ± 0.08 | 3.27 ± 0.26 | 2.90 ± 0.09 | 2.48 ± 0.26 | 2.86 ± 0.14 | 2.59 ± 0.15 | 1.90 ± 0.18 |
| $a$ | 0.66 ± 0.02 | 0.42 ± 0.02 | 0.55 ± 0.01 | 0.52 ± 0.04 | 0.47 ± 0.01 | 0.50 ± 0.01 | 0.60 ± 0.02 |
| $r_{inf}$ | 0.007±0.001 | 0.019±0.003 | 0.031±0.001 | 0.012±0.002 | 0.015±0.003 | 0.011±0.001 | 0.006±0.002 |
| **5P** | | | | | | | |
| $\rho_{dye}$ | 0.53 ± 0.04 | 0.56 ± 0.03 | 0.34 ± 0.01 | 0.55 ± 0.03 | 0.55 ± 0.04 | 0.57 ± 0.02 | 0.52 ± 0.02 |
| $\rho_{segment}$ | 1.75 ± 0.13 | 2.56 ± 0.15 | 2.98 ± 0.10 | 3.13 ± 0.07 | 2.35 ± 0.14 | 2.01 ± 0.05 | 1.43 ± 0.08 |
| $a$ | 0.72 ± 0.03 | 0.51 ± 0.23 | 0.59 ± 0.01 | 0.41 ± 0.18 | 0.54 ± 0.21 | 0.51 ± 0.01 | 0.61 ± 0.03 |
| $r_{inf}$ | 0.001±0.001 | 0.017± 0.007 | 0.027± 0.001 | 0.021± 0.009 | 0.013± 0.006 | 0.010± 0.001 | 0.004± 0.001 |
| **NP + 4E** | | | | | | | |
| $\rho_{dye}$ | 0.52 ± 0.01 | 0.62 ± 0.03 | | 0.62 ± 0.05 | 0.61 ± 0.03 | 0.65 ± 0.04 | 0.59 ± 0.02 |
| $\rho_{segment}$ | 1.93 ± 0.13 | 3.36 ± 0.11 | | 2.65 ± 0.13 | 4.54 ± 0.21 | 5.35 ± 0.45 | 3.34 ± 0.27 |
| $a$ | 0.70 ± 0.02 | 0.49 ± 0.01 | | 0.51 ± 0.04 | 0.46 ± 0.01 | 0.43 ± 0.01 | 0.59 ± 0.02 |
| $r_{inf}$ | 0.020±0.001 | 0.045± 0.001 | | 0.030± 0.004 | 0.075± 0.004 | 0.077± 0.009 | 0.043± 0.002 |
| **NP DEN*** | | | | | | | |
| $\rho_{dye}$ | 0.60 ± 0.06 | 0.53 ± 0.03 | | 0.52 ± 0.03 | 0.51 ± 0.05 | 0.51 ± 0.04 | 0.53 ± 0.03 |
| $\rho_{segment}$ | 0.89 ± 0.10 | 1.54 ± 0.10 | | 1.31 ± 0.04 | 1.43 ± 0.08 | 1.42 ± 0.07 | 0.95 ± 0.06 |
| $a$ | 0.53 ± 0.11 | 0.39 ± 0.03 | | 0.42 ± 0.02 | 0.41 ± 0.05 | 0.40 ± 0.04 | 0.42 ± 0.03 |
| $r_{inf}$ | 0.001±0.001 | 0.012± 0.001 | | 0.006± 0.002 | 0.008± 0.002 | 0.012± 0.001 | 0.006± 0.001 |
| **5P DEN*** | | | | | | | |
| $\rho_{dye}$ | 0.51 ± 0.05 | 0.47 ± 0.01 | | 0.49 ± 0.04 | 0.50 ± 0.08 | 0.53 ± 0.03 | 0.46 ± 0.09 |
| $\rho_{segment}$ | 0.82 ± 0.08 | 1.27 ± 0.04 | | 1.57 ± 0.05 | 1.43 ± 0.19 | 1.49 ± 0.09 | 0.96 ± 0.09 |
| $a$ | 0.42 ± 0.09 | 0.38 ± 0.01 | | 0.39 ± 0.03 | 0.42 ± 0.07 | 0.41 ± 0.02 | 0.42 ± 0.09 |
| $r_{inf}$ | 0.004±0.002 | 0.010± 0.003 | | 0.012± 0.003 | 0.015± 0.003 | 0.008± 0.001 | 0.004± 0.001 |

[a] All data were fitted to **Eq. 1**. Rotational correlation lifetimes ($\rho$) are given in nanoseconds. Error values were derived as described in the Methods section in the main text. *DEN refers to samples in a solution containing 6M GdmCl, rotational correlation times in 6M GdmCl were viscosity corrected.

**Table S3.** FCS decay parameters (lifetimes and amplitudes) for different states of 4E-BP2 labelled at different sites along the sequence [a].

| | C0 | S14C | C35 | C73 | S91C | C121 |
|---|---|---|---|---|---|---|
| **NP** | | | | | | |
| $\tau_1$ | 57.1 ± 3.9 *(0.13 ± 0.01)* | 59.7 ± 4.0 *(0.13 ± 0.01)* | 53.7 ± 4.6 *(0.11 ± 0.01)* | 55.5 ± 6.2 *(0.11 ± 0.01)* | 67.2 ± 3.4 *(0.17 ± 0.01)* | 54.3 ± 4.0 *(0.12 ± 0.01)* |
| $\tau_t$ | 8.8 ± 1.1 *(0.06 ± 0.01)* | | | | | |
| $\tau_2$ | - | - | 0.31 ± 0.07 *(0.11 ± 0.01)* | 0.20 ± 0.09 *(0.11 ± 0.01)* | - | - |
| $\tau_D$ | 211 ± 1 | | | | | |
| **5P** | | | | | | |
| $\tau_1$ | 83.9 ± 2.2 *(0.23 ± 0.01)* | 70.1 ± 2.1 *(0.17 ± 0.01)* | 79.4 ± 2.6 *(0.18 ± 0.01)* | 64.9 ± 2.5 *(0.14 ± 0.01)* | 65.2 ± 2.0 *(0.16 ± 0.01)* | 71.7 ± 2.7 *(0.16 ± 0.01)* |
| $\tau_t$ | 7.5 ± 0.8 *(0.04 ± 0.01)* | | | | | |
| $\tau_2$ | - | - | - | - | - | - |
| $\tau_D$ | 212 ± 1 | | | | | |
| **NP + 4E** | | | | | | |
| $\tau_1$ | 96.3 ± 1.9 *(0.28 ± 0.01)* | 101.8 ± 2.6 *(0.22 ± 0.01)* | 100.5 ± 2.1 *(0.25 ± 0.01)* | 99.9 ± 2.9 *(0.19 ± 0.01)* | 101.0 ± 2.0 *(0.26 ± 0.01)* | 59.2 ± 8.0 *(0.15 ± 0.01)* |
| $\tau_t$ | 10.0 ± 0.8 *(0.06 ± 0.01)* | | | | | |
| $\tau_2$ | - | - | 0.10 ± 0.04 *(0.16 ± 0.05)* | 0.04 ± 0.01 *(0.35 ± 0.11)* | 0.14 ± 0.03 *(0.19 ± 0.03)* | 0.14 ± 0.02 *(0.16 ± 0.01)* |
| $\tau_D$ | 289 ± 1 | | | | | |

[a] All data were fitted to **Eq. 2** Lifetimes are given in microseconds, with corresponding amplitudes between brackets. Error values were derived as described in the Methods section in the main text.

**Table S4.** Absolute solvent accessible surface area (SASA) and relative surface area (RSA) for eIF4E (PDB: 3AM7) tryptophan and tyrosine residues, calculated using the POPS algorithm(13).

| **Residue** | **W43** | **W46** | **W56** | **W73** | **W102** | **W113** | **W130** | **W166** | **Y34** | **Y76** | **Y91** | **Y145** | **Y183** | **Y197** |
|---|---|---|---|---|---|---|---|---|---|---|---|---|---|---|
| **SASA (Å$^2$)** | 18.5 | 46.3 | 87.9 | 63.5 | 125.2 | 20.5 | 28.1 | 27.5 | 82.8 | 51.4 | 30.4 | 44.4 | 25.5 | 11.1 |
| **RSA** | 0.094 | 0.24 | 0.45 | 0.32 | 0.64 | 0.10 | 0.14 | 0.14 | 0.40 | 0.25 | 0.15 | 0.22 | 0.12 | 0.054 |
